# CIP2A interacts with TopBP1 and is selectively essential for DNA damage-induced basal-like breast cancer tumorigenesis

**DOI:** 10.1101/2020.08.27.269902

**Authors:** Anni Laine, Srikar G. Nagelli, Caroline Farrington, Umar Butt, Anna N. Cvrljevic, Julia P. Vainonen, Femke M. Feringa, Tove J. Grönroos, Prson Gautam, Sofia Khan, Harri Sihto, Xi Qiao, Karolina Pavic, Denise C. Connolly, Pauliina Kronqvist, Laura L. Elo, Jochen Maurer, Krister Wennerberg, Rene H. Medema, Heikki Joensuu, Emilia Peuhu, Karin de Visser, Goutham Narla, Jukka Westermarck

## Abstract

Despite saturated genetic profiling of breast cancers, oncogenic drivers for the clinically challenging basal-like breast cancer (BLBC) subtype are still poorly understood. Here, we demonstrate that CIP2A is selectively essential for DNA damage-induced initiation of mouse BLBC tumors, but not of other cancer types. Mechanistically, CIP2A was discovered genome-widely the closest functional homologue for DNA-damage proteins TopBP1, RHNO, POLQ, NBN and PARP1. CIP2A directly interacts with the ATR-activation domain of TopBP1, and dampens both, chromatin binding of TopBP1 and RAD51, and G2/M checkpoint in DNA-damaged cells. CIP2A also drives BLBC-associated proliferative MYC and E2F1 signaling. Consistently with high DNA-damage response activity BLBCs, and CIP2A’s novel role in checkpoint signaling, CIP2A was found essential for DNA-damaged, and BRCA-mutant BLBC cells. Selective role for CIP2A as BLBC driver was supported by association of high CIP2A expression with poor patient prognosis only in BLBC, but not in other breast cancer types. Therapeutically, small molecule reactivators of PP2A (SMAPs) phenocopy CIP2A-dependent DNA damage response, and inhibit *in vivo* growth of patient-derived BLBC xenograft. In summary, we discover sub-type selective essential role for CIP2A in BLBC initiation and maintenance that can be explained by its newly discovered association with DNA-damage response, coordinated with regulation of proliferative signaling. The results also identify therapeutic strategy for CIP2A-dependent BLBCs.

## Introduction

Breast cancer is classified into molecular subtypes based on their cell surface receptor expression and transcriptional profiles. One of the most aggressive and clinically challenging breast cancer subtype is the basal-like breast cancer (BLBC)^1–3^. The hallmarks of BLBCs are high genetic instability, BRCA mutations, TP53 inactivation, constitutive DNA damage response (DDR) signaling, dysregulation of EGFR, and high proliferation activity ^1–3^. About 75% of BLBCs belong to the triple-negative breast cancer subtype (BL-TNBCs), devoid of ER, PR and HER2 ^1^. In addition to their frequently aggressive clinical appearance, the lack of these targetable receptors makes BLBCs therapeutically very challenging. Therefore, characterization of oncogenic driver(s) responsible for BLBC initiation and disease progression could provide novel opportunities for BLBC therapy.

Despite the near saturated genetic knowledge of breast cancer, no clear genetic oncogenic drivers have been identified for the BLBCs ^1, 3^. This indicates that BLBC is radically different from other breast cancer subtypes driven by either receptor tyrosine kinase activity in the case of HER2 positive breast cancers, or by hormonal receptor-mediated transcriptional programs such as in ER and PR positive breast cancers. The high proliferation activity in BLBCs can be accounted to loss of cell cycle inhibition by p53 mutations and to high EGFR activity, but there is no evidence that they alone, or in combination, would be sufficient for tumor initiation in BLBC. Beside high proliferation activity, genomic instability and high DDR activity are important hallmarks of BLBC ^2, 3^. Most clinical BLBCs are also deficient for homologous recombination (HR), either through the acquisition of BRCA mutations or other defects in the HR pathways. Based on these hallmarks of BLBC, it could be hypothesized that potential drivers of this breast cancer subtype has to both support high proliferation activity, and also dampen the cell cycle effects of tumor suppressive DDR activity ^4, 5^.

Healthy cells respond to double stranded DNA breaks (DSB) by activation of the G2/M cell cycle checkpoint and consequent mitotic arrest ^5^. To allow mitotic progression under DNA damaging conditions, transformed cells instead have developed (phosphorylation-dependent) strategies to dampen G2/M checkpoint signaling ^5, 6^. These mechanisms are important in the early phases of tumor initiation by allowing the mitotic progression of DNA damaged premalignant cells. One of the DDR proteins involved in G2/M checkpoint signaling is DNA Topisomerase II binding protein 1 (TopBP1)^7, 8^, which is a scaffold protein interacting with checkpoint kinase ATR through its ATR-activation domain (AAD)^9^. In the presence of DSBs, TopBP1 promotes RAD51 chromatin loading resulting in G2/M arrest ^10–14^. These features make TopBP1 an interesting effector for G2/M checkpoint dampening in cancer cells ^5, 8, 12^, but to date its regulation and importance in BLBC cells has remained largely unknown.

While kinase dysregulation appears to be insufficient to drive BLBC initiation, the role of their counterparts, phosphatases remain to be poorly understood. Recently serine/threonine phosphatase PP2A have gained attention as a druggable tumor suppressor ^15–17^. Especially the role of serine/threonine phosphatases in DNA damage response at chromatin ^6^, could link them to cancer types with high mutation burden such as BLBCs. PP2A is inhibited in most cancers by non-genetic mechanisms including high expression of endogenous inhibitor proteins such as CIP2A, PME-1 or SET^17, 18^. *CIP2A* gene is not genetically prevalently mutated in any cancer type (https://cancer.sanger.ac.uk/cosmic), and it is only expressed at low levels in normal mammary gland tissue. However, *CIP2A* transcription is induced by *TP53* mutation via E2F1 activity ^19, 20^, and by EGFR ^21, 22^, all features closely linked to BLBC. However, it is currently unclear what is CIP2A’s potential role in BLBC initiation, maintenance, or therapeutic targeting. In general, it is unclear whether CIP2A, or any of the PP2A inhibitor proteins, are essential for initiation of any cancer type? Notably, understanding of CIP2A-related cancer initiation mechanisms is also therapeutically relevant due to recent development of small molecule activators of the CIP2A-inhibited PP2A-B56 with potent antitumor activities in several preclinical cancer models *in vivo* ^15, 16^.

In this study, we provide first evidence for essential role for PP2A inhibitor protein in tumor initiation. Specifically, we demonstrate that CIP2A is selectively essential for initiation of BLBC, but not of other mouse tumor types. Further, among transformed breast cancer cell types, CIP2A is selectively essential for survival of BRCA/TP53-mutant BLBC cells. Mechanistically this can be explained by previously unidentified profound functional similarity between CIP2A and the core DDR proteins; and subsequent role for CIP2A in preventing RAD51 recruitment to chromatin upon DNA-damage. CIP2A also promotes MYC and E2F1 activities in BLBC cells; Finally, we discover that SMAPs transcriptionally inhibit *CIP2A* expression and serve as candidate therapeutics for CIP2A-positive BLBCs.

## Results

### *Cip2a* is selectively required for initiation of DMBA-induced mammary tumors in mice

Thus far the only evidence for the importance of CIP2A for *in vivo* tumor initiation is modest reduction of number of HER2-driven mammary tumors in the genetic crosses between transgenic MMTV-neu and *Cip2a*-deficient (*Cip2a*−/−) mouse models ^20^. Thus, it is yet totally unclear whether CIP2A is essential for initiation of any cancer (sub)type *in vivo.* To address this question, we challenged the previously described *Cip2a*−/− mice ^20, 23^ with a chemical carcinogenesis protocol consisting only of six consecutive doses of the genotoxic agent 7,12-dimethylbenz[a]anthracene (DMBA)(Fig. 1A). Similar to other polyaromatic hydrocarbons, DMBA forms covalent DNA adducts, and induces a DNA damage response (DDR) including activation of ɣH2AX, ATR, and RAD51 ^24, 25^. Oral exposure of mice with DMBA induces mouse BLBCs ^26^, but also several other cancers ^27^, allowing us to assess the relative importance of *Cip2a* across different mouse cancer types. Importantly, as compared to models combining DMBA and hormones, such as progestin Medroxyprogesterone Acetate (MPA), the DMBA-only mammary tumors are initiated with much longer latency^28^, better resembling course of human breast cancer development. Molecularly DMBA-induced BLBCs are also different form *Brca*/*p53* mutant, or transgenic Wnt-induced tumors. For example, whereas deletion of either *Brca1* or *Brca2* abrogates Rad51 recruitment upon DNA-damage ^29^, basal cells from DMBA model have retained this DDR mechanism relevant to cell cycle arrest in S-phase^10–12, 26^. Thereby use of DMBA-induced model, in which the tumor initiating cell population is basal cells ^28^, could allow discovery of BLBC driver mechanisms not necessarily revealed by the other models.

**Figure 1:**
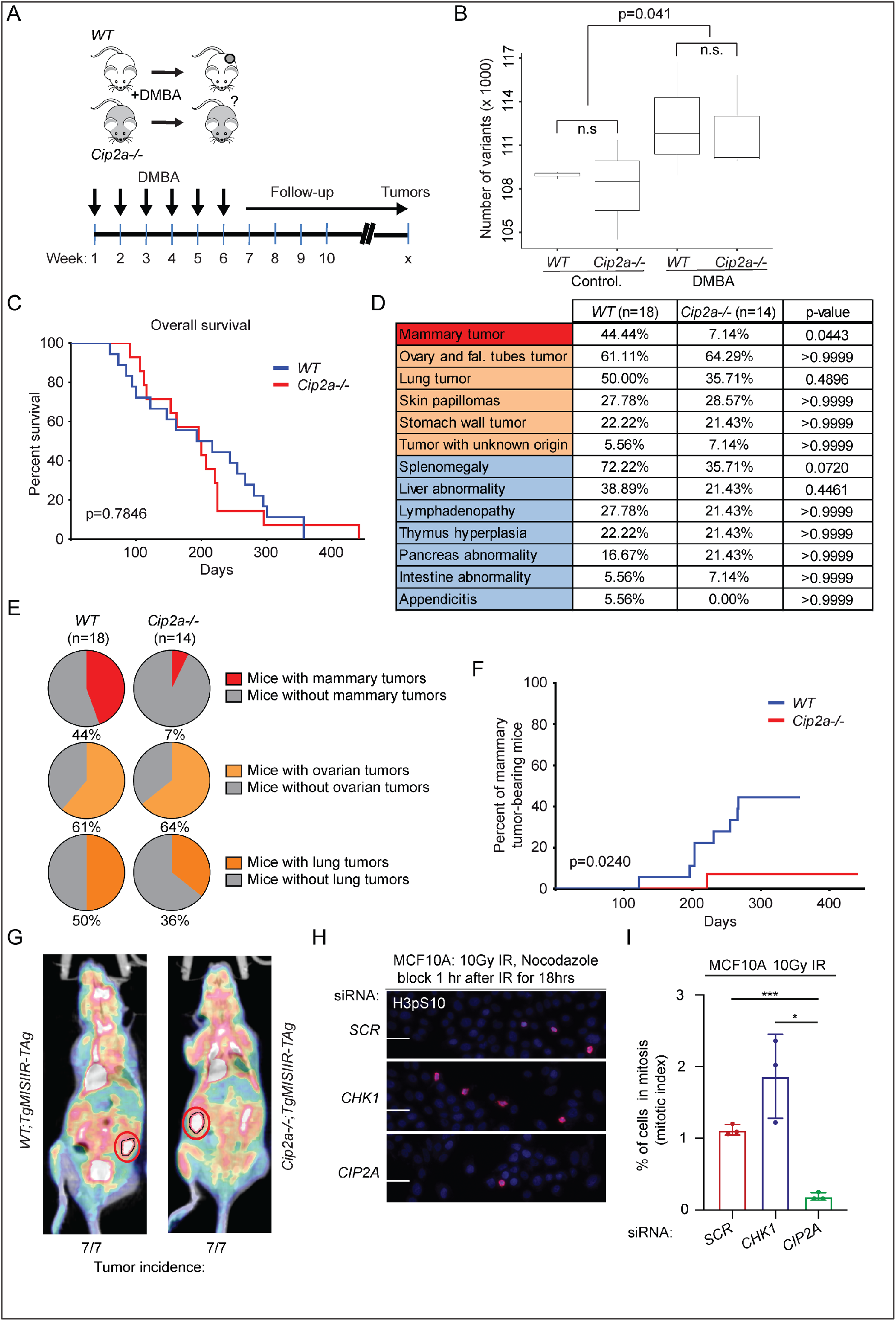
*Cip2a* knockout mice are selectively resistant to DMBA-induced mammary tumorigenesis. **A**, A schematic presentation of the chemical *in vivo* carcinogenesis mouse model. DMBA was orally administered to wild type (*WT*) and *Cip2a*−/− mice once a week for 6 consecutive weeks after which mice were monitored for signs of spontaneous tumor formation. **B**, Number of genetic variants in exons of the expressed genes in non-treated (control) and DMBA-treated *WT* (n=3) and *Cip2a*−/− (n=3) mouse mammary glands. P-value by Wilcoxon test. **C**, Overall survival of *WT* and *Cip2a*−/− mice in days after starting of DMBA administration. Shown is the survival of 18 *WT* and 14 *Cip2a*−/− mice. P-value by log-rank test. **D**, Incidences of tumor formation in different tissues and other pathologies in sacrificed DMBA-administered *WT* (n=18) and *Cip2a*−/− (n=14) mice. P-values between *WT* and *Cip2a*−/− groups in each pathology calculated by Fisher’s exact test. **E**, Proportion of the most common tumor types induced by DMBA in *WT* and *Cip2a*−/− mice. Percentage of tumor carrying mice in both groups shown under pie charts. **F**, Incidence of mammary tumors in *WT* (n=18) and *Cip2a*−/− (n=14) mice presented in days after starting administration of DMBA. P-value by log-rank test. **G**, Incidence of ovarian tumors in *WT;TgMISIIRTAg* and *Cip2a*−/−;*TgMISIIR-Tag* mice. Mice imaged by metabolic active tumor volume (MATV) definition by ^18^F-FDG PET/CT imaging. Red circle denotes the tumor. **H**, Mitotic index analysis of MCF10A cells transfected with the indicated siRNAs. *CHK1* siRNA was used as positive control. Cells were treated with 10Gy radiation dose and Nocodazole (100 ng/ml) block 1 hour after IR for 18 hours. Mitotic cells were stained using phospho-histone H3 at Ser10. Scale bar: 100μm. **I**, % of H3pS10 positive nuclei in pooled form from n=3 replicates, expressed as mean ± SD.

As expected ^25^, DMBA treatment induced a significant increase in mutation load in non-tumorigenic mammary gland tissue already 2 weeks after the last DMBA dosing; however the mutation load (Fig. 1B), or overall survival was not associated with *Cip2a* genotype (Fig. 1C). When assessed by palpation, external observation, and by tissue pathology analysis upon autopsy of the mice with any symptoms of reduced well-being, tumors in five different tissue types were observed in the DMBA-treated mice (Fig. 1D). In addition, mice displayed other pathological phenotypes mostly associated with lymphadenopathy. Notably, while incidence of tumors in ovary, lung, skin or stomach were not altered in *Cip2a*−/− mice, mammary tumors showed almost absolute dependence on *Cip2a* for tumor initiation (Fig. 1D-F). To control that lack of genotype dependence of other cancer types on *Cip2a* was not due to leakage of genetrap cassette used for *Cip2a* gene silencing ^23^, we confirmed the absence of CIP2A protein expression in ovarian cancer tissues from *Cip2a*−/− mice (Fig. S1A). We further confirmed that *Cip2a* was dispensable for skin and ovarian tumorigenesis by independent *in vivo* models. To this end, we crossed *Cip2a*−/− mice with the MISIIR-Tag mouse model producing tumors resembling high grade ovarian cancer ^30^, but did not observe any notable difference in ovarian tumorigenesis between *Cip2a* wild-type (*WT*) or *Cip2a*−/− mice by PET/CT-imaging or by visual inspection after autopsy (Fig. 1G, S1B). For the skin tumorigenesis we used classical DMBA/TPA two-stage skin tumorigenesis protocol as described in the supplementary materials and methods. Nevertheless, there was no difference in skin tumor initiation between *Cip2a* genotypes (Fig. S1C).

Results above strongly indicate that CIP2A is required for propagation of DNA-damaged mammary epithelial cells. To validate that this is a cell intrinsic property of CIP2A, we tested the impact of CIP2A silencing on mitotic progression of MCF-10A basal like immortalized mammary epithelial cells treated with ionizing radiation (IR). Notably, whereas inhibition of checkpoint kinase CHK1 abrogated G2/M checkpoint, and CIP2A silencing did not impact mitotic progression of untreated MCF-10A cells (Fig. S1D), CIP2A was found indispensable for G2/M progression in IR-treated MCF10A cells (Fig. 1H). To provide independent validation to these results, and to assess the selectivity of CIP2A among other PP2A inhibitor proteins, we surveyed results from a genetic screen in HAP1 cells ^31^(see Fig. S1E for technical description). Directly supportive of the results in MCF-10A cells, CIP2A was the only tested PP2A inhibitor protein that became significantly essential under repeated low-dose irradiation (Fig. S1F).

These results establish notable selective essentiality for *Cip2a* for mitotic progession of DNA-damaged cells, and for the initiation of DNA-damage induced mammary tumors previously defined to represent mouse BLBCs^26^. As such the results represent first evidence for potential cancer driver role for CIP2A in any cancer type.

### *Cip2a* is induced by DMBA in premalignant mammary gland tissue and drives initiation of mouse BLBC-like tumors

A key criterion for a cancer driver candidate involved in tumor initiation, is expression in premalignant tissue prior tumorigenesis. To examine this, we studied *Cip2a* mRNA expression in non-tumorigenic mammary gland tissues from control and DMBA-treated animals, and from DMBA-induced mammary tumors in *Cip2a WT* mice. Consistent with negligible CIP2A protein expression in normal human mammary glands ^28^, *Cip2a* mRNA was expressed at a very low level in control mouse mammary glands (Fig. 2A). Importantly, mammary glands sampled 2 weeks after the 6th dose of DMBA (Fig. 1A) displayed significantly increased *Cip2a* mRNA expression (Fig. 2A). In line with suggested role as a disease driver, *Cip2a* mRNA expression was induced significantly further in mammary tumors from DMBA-treated *WT* mice (Fig. 2A).

**Figure 2:**
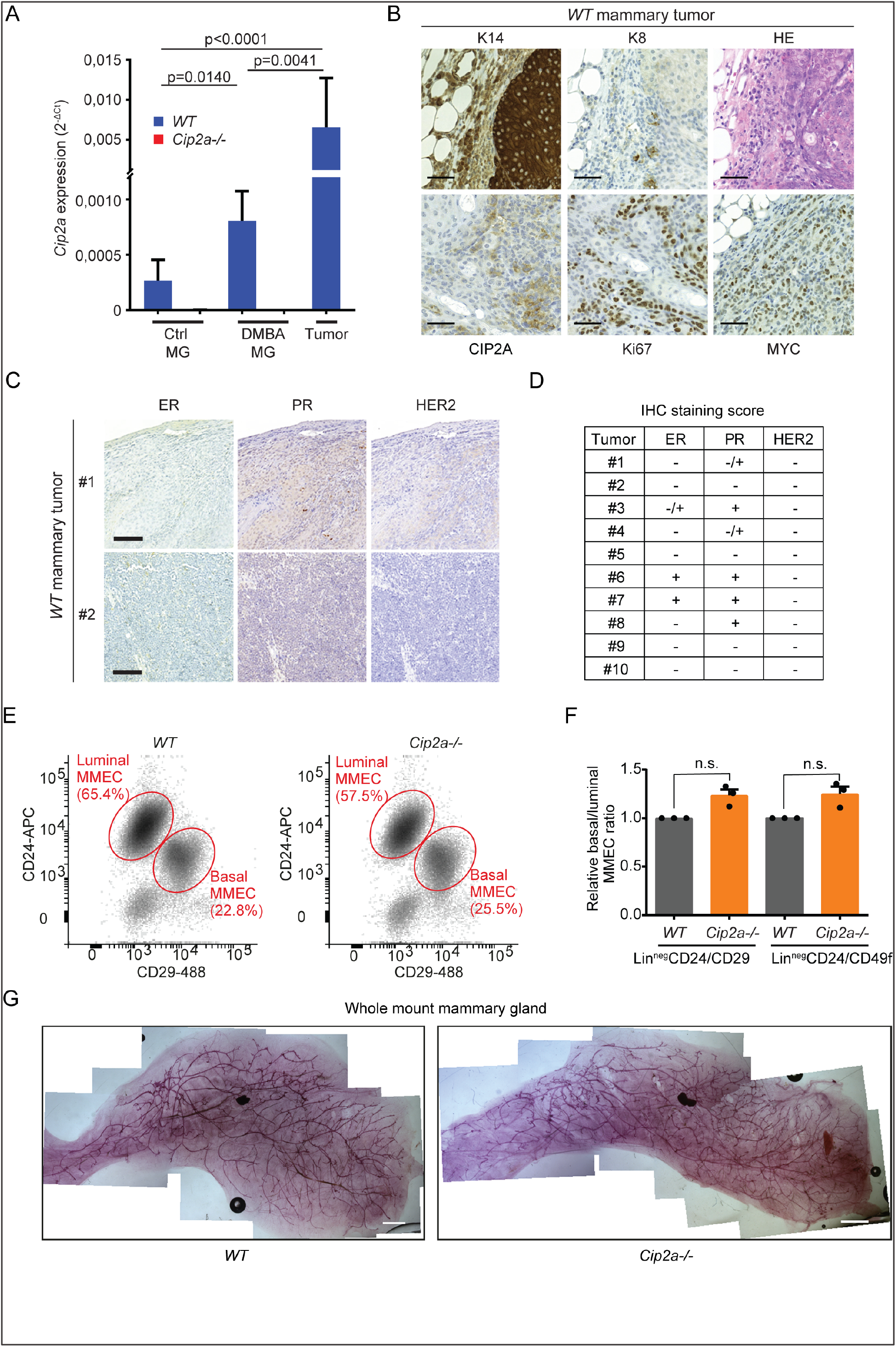
*Cip2a* drives initiation of mouse BLBC-like tumors but is dispensable for normal mammary gland development. **A**, qRT-PCR analysis of *Cip2a* mRNA expression normalized to *Actb* and *Gapdh* from *WT* and *Cip2a*−/− non-treated (Ctrl) and DMBA-administered mouse non-tumorigenic mammary glands (MG), and WT DMBA-induced mammary tumors. Shown is mean ± SD of 10 *WT* and 9 *Cip2a*−/− non-treated mammary glands (Ctrl MG), 3 *WT*, and 3 *Cip2a*−/− mammary glands from DMBA-administered mice, and 16 mammary tumors from *WT* DMBA-induced mice. P-values calculated by Mann-Whitney test. **B**, Immunohistochemical characterization of DMBA-induced mammary tumors from *WT* mice. Shown are representative images of immunohistochemical staining of Keratin-14 (K14), Keratin-8 (K8), CIP2A, Ki67 and MYC proteins and hematoxylin and eosin (HE) histochemical staining. Scale bar: 50μM **C**, Immunohistochemical characterization of DMBA-induced mammary tumors from *WT* mice for receptor status. Shown are representative images of immunohistochemical staining of estrogen receptor (ER), progesterone receptor (PR) and HER2 from two individual tumors. Scale bar: 100μM **D**, Semiquantitative analysis of receptor status from 10 individual WT tumors. **E**, Mouse mammary epithelial cells (MMECs) isolated from *WT* and *Cip2a*−/− mice were immunolabelled for surface markers. Among the lineage-negative cells (CD31^neg^, CD45^neg^), the basal epithelial (CD24^low-neg^, CD29^high^) and luminal epithelial (CD24^pos^, CD29^low-neg^) cell populations were quantified by flow cytometry. The gates and % of cells are indicated in red. **F**, The ratio between basal and luminal epithelial cells in each sample calculated using two different labelling strategies (CD24/CD29 and CD24/CD49f) and pooled from independent experiments (n=3). Data are mean ± SEM. P-values calculated by unpaired t-test. **G**, Representative images of mammary gland whole mounts from adult *WT* and *Cip2a*−/− mice. Scale bar: 2 mm.

Next, we conducted molecular characterization of the mammary tumors from DMBA-treated *WT* mice. Consistent with a previous report demonstrating that the tumor initiating cells from the DMBA model are of basaloid origin^26^, we observed a BLBC and BL-TNBC phenotypes in majority of the characterized tumors (Fig. 2B-D). Also consistent with BLBC phenotype, the tumors in *WT* mice were highly proliferative based on Ki67 staining, and displayed MYC protein overexpression (Fig. 2B). Notably, the lack of predominantly BLBC tumors in *Cip2a*−/− mice was not related to any genotype-associated alterations in the basal and luminal epithelial cell ratio in the mammary gland (Fig. 2E,F). The purity of basal and luminal fractions was assessed by qRT-PCR (Fig. S2A,B). Furthermore, consistent with the very low expression of *Cip2a* in normal mammary glands (Fig. 2A), and the normal nursing behavior of the *Cip2a*−/− mice, we did not observe any notable differences in the mammary gland development and branching morphogenesis between *WT* and *Cip2a*−/− mice (Fig. 2G). Collectively, these results demonstrate that although *Cip2a* is dispensable for normal mouse mammary development, DNA-damage-elicited induction of *Cip2a* mRNA expression is required for initiation of mouse BLBC-like tumors.

### Co-dependence analysis reveals a functional association between CIP2A, TopBP1 and homology directed DNA repair

CIP2A promotes several different oncogenic mechanisms across human cancer types ^20, 32, 33^. However, as the currently known mechanisms regulated by CIP2A do not explain the selective essentiality of CIP2A for DNA-damage-induced BLBC initiation (Fig. 1), we hypothesized that CIP2A promotes BLBC initiation by yet uncharacterized DNA-damage associated mechanism. To identify such mechanism, we surveyed a CRISPR/Cas9-based dropout screen repository from DepMap (Avana 2020 Q1; https://depmap.org), to identify genes in an unbiased manner that are most significantly similar in their essentiality with *CIP2A* across all 739 human cancer cell lines. Consistent with the observed *Cip2a*-dependency of DNA-damage-induced tumorigenesis (Fig. 1), the top 10 co-dependent genes with *CIP2A* (i.e. functionally most similar to CIP2A) were all associated with DNA repair (Fig. 3A). Notably, out of the top ten *CIP2A*-associated DNA repair factors, *CIP2A* was at the genome-wide level the most significantly similar gene for *RHNO1, TOPBP1*, *POLQ, NBN* and *PARP1* (Fig. 3A,B). In the case of *TOPBP1*, the co-dependency with *CIP2A* was greater than with *ATR* (Fig. 3B), which is the bona-fide TopBP1 DDR effector ^8, 9^. Although surprising, these results are supported by recent screening results implicating essentiality of both CIP2A and TopBP1 for recovery of cancer cells from ATR inhibition ^34^. When analyzed for functional protein association networks by STRING database (https://string-db.org), the CIP2A-associated proteins (Fig. 3A) formed a tight protein network (Fig. 3C) that was functionally linked with processes such as “Homology directed Repair”, “G2/M DNA damage checkpoint”, and “Processing of DNA double-strand break ends” (Fig. 3D). Interestingly, in a recent PP2A-related phosphoproteome survey ^35^, CIP2A was found to prevent the dephosphorylation of Nibrin (NBN) which was one of the TopBP1 protein network members and is known to co-operate with TopBP1 in ATR activation ^36^(Fig. 3C). Additional evidence for the intertwining of CIP2A with the TopBP1 complex, and mitosis, was obtained by mRNA co-expression analysis across 1156 cell lines from the Broad institute Cancer Cell line Encyclopedia ^37^(Fig. S3A). Reactome pathway analysis of the 10 genes most significantly co-expressed with *CIP2A* revealed clear enrichment of mitotic genes (Fig. 3E). Of the *CIP2A* co-dependent genes (Fig. 3A), *TOPBP1* and *POLQ* were also among the 25 most significantly co-expressed genes with CIP2A (Fig. 3F and Fig. S3A). Both these genes showed also very significant co-expression with *CIP2A* in BLBC (Fig. S3B). Collectively these results reveal an intimate, but previously unidentified association of CIP2A with critical DNA repair complex proteins, and with homology directed DNA repair in mitosis; potentially highly relevant to the role of CIP2A in facilitating malignant progression towards BLBC under DNA damaging conditions *in vivo* (Fig. 1).

**Figure 3:**
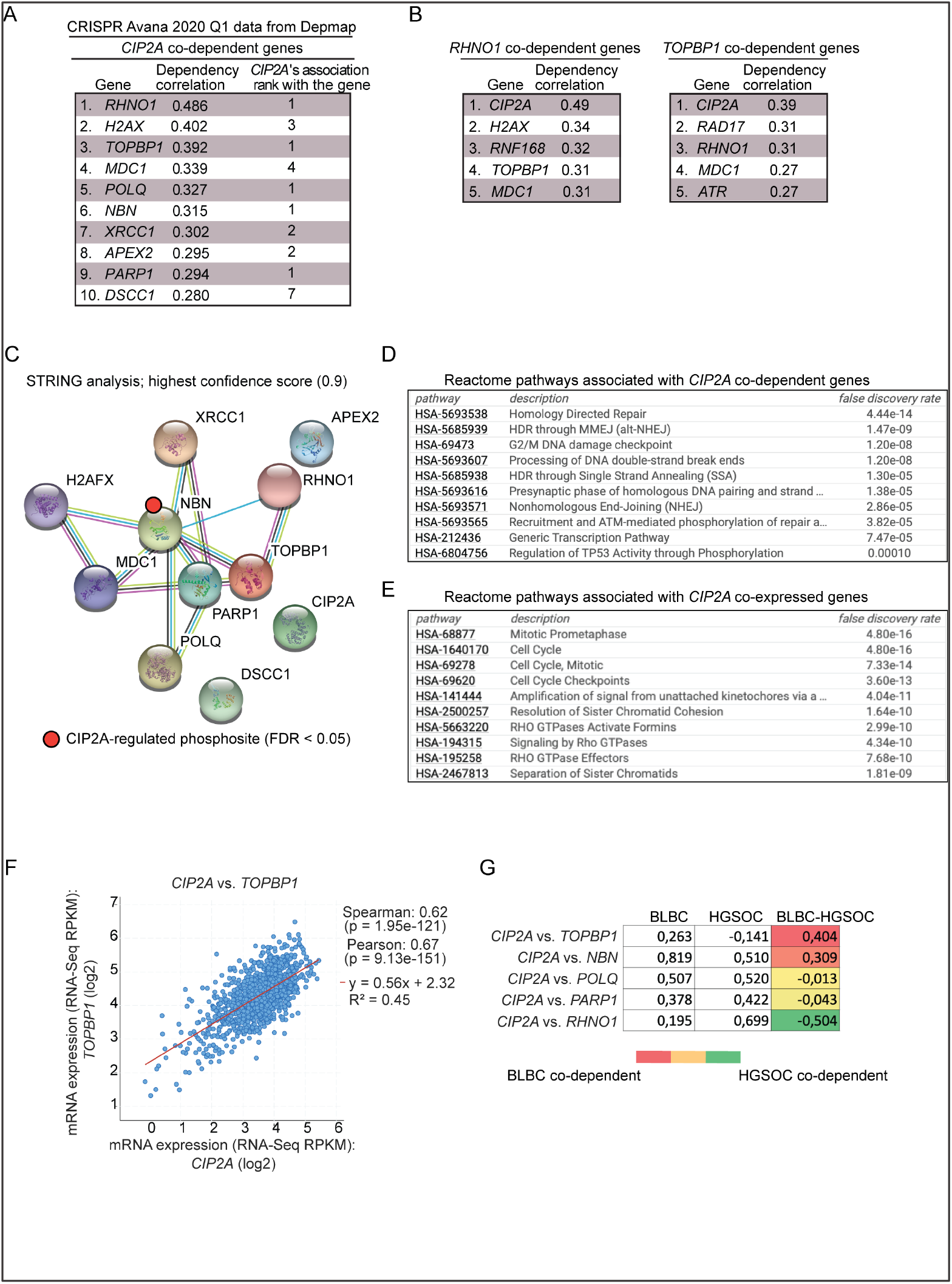
Co-dependence analysis reveals functional association of CIP2A with critical DNA damage response proteins. **A**, Top 10 co-dependencies of *CIP2A* across 739 cell lines genome-wide from CRISPR Avana screen. *CIP2A*’s own co-dependency rank for the top 10 genes is also listed. Data extracted from DepMap portal (Avana 2020Q1). **B**, Genome-widely, *CIP2A* is the closest functional homologue to *RHNO1* and *TOPBP1*. **C**, STRING functional protein association network analysis of CIP2A co-dependent proteins from (A). By using the highest data confidence score (0.9), except for APEX2, DSCC1 and CIP2A, the other proteins form highly connected protein network. NBN phosphorylation indicated by red dot was found to be regulated by CIP2A based on ^35^. **D**, Top 10 Reactome pathways associated with genes from (A). **E**, Top 10 Reactome pathways associated with CIP2A co-expressed genes derived from Cancer Cell Line Encyclopedia (1156 samples). **F**, Correlation between *CIP2A* and *TOPBP1* mRNA expression across 1156 cell lines from Cancer Cell Line Encyclopedia. **G**, Pair-wise correlation of dependence of either BLBC or HGSOC cell lines of the indicated genes from DepMap portal (Avana 2020Q1). The values for BLBC and HGSOC indicates correlation (max. 1) in dependence of the cells for the genes in the gene pair; the higher number indicating for higher similarity in the dependence. The color-coded numbers indicate the difference in the co-dependence between BLBC and HGSOC cells for the indicated gene pair.

The DepMap co-dependence data was also utilized to understand the interesting difference in CIP2A dependence in the initiation of mammary and ovarian cancers, as BLBC and high grade serous ovarian cancer (HGSOC) are known to share similar characteristics. To this end, we analyzed in a pair-wise fashion the correlation between dependence on either *CIP2A*, or one of the genes *RHNO1, TOPBP1*, *POLQ, NBN* and *PARP1* across either BLBC or HGSOC cell lines. In BLBC, *TOPBP1* and *NBN* had higher co-dependence with CIP2A than in HGSOC cells, while in HGSOC, *RHNO1* was more co-dependent with *CIP2A* (Fig. 3G). These differences may provide one plausible explanation for the differential requirement of *Cip2a* for DMBA-induced BLBC-like, but not ovarian cancer initiation (Fig. 1D,E,G). Notably, *TOPBP1* was the only studied gene which did not show *CIP2A* co-dependence in HGSOC cells, but was co-dependent in BLBC cells (Fig. 3G), strengthening the role of TopBP1 as the candidate mechanistic link between CIP2A, and malignant progression of DNA-damaged BLBC cells.

### CIP2A dampens TopBP1-RAD51 function under DNA damage

The results above identify CIP2A as a novel candidate protein involved in the function of TopBP1 in double stranded DNA damage repair, and in G2/M arrest. However, as illustrated by the STRING analysis (Fig. 3C), there is currently no evidence for direct mechanistic link between CIP2A and the TopBP1 complex. Here, by using a genome-wide Y2H assay with human breast cancer cDNA library, TopBP1 was identified with very high confidence as a direct interaction partner for CIP2A (Table S1 and Fig. 4A). Interaction between TopBP1 and endogenous nuclear CIP2A ^38^ was confirmed by co-immunoprecipitation analysis (Fig. 4B), and by proximity ligation analysis (Fig. S4A). The interaction with CIP2A was delineated to be mediated by the 6^th^ BRCT domain of TopBP1, both by matching the interacting regions from overlapping TopBP1 fragments in the Y2H assay (Fig. 4A, Table S1), and by co-immunoprecipitation analysis (Fig. 4C,D). Notably, the interaction was greatly strengthened by the presence of the ATR-activation domain (AAD) of TopBP1 adjacent to 6^th^ BRCT repeat (Fig. 4C,D). Functionally, removal of CIP2A resulted in constitutive ATR activation in the basal-type premalignant mammary cell line MCF-10A (Fig. 4E). As a more direct evidence linking CIP2A to TopBP1-regulated DDR, the highest H2AX phosphorylation (γH2AX) was observed in CIP2A-depleted cells overexpressing TopBP1 variant that contains AAD (Fig. 4F). γH2AX also co-immunoprecipitated with TopBP1 and CIP2A from DNAse treated cellular lysates (Fig. 4B). As further support for the role of CIP2A in dampening TopBP1 function, CIP2A depletion resulted in enhanced chromatin recruitment of TopBP1 in X ray-irradiated (IR) MCF10A cells (Fig. 4G,H). This was specific to TopBP1, as CIP2A did not impact IR-induced p53BP1 chromatin recruitment (Fig. S4B). Furthermore, consistently with the role of both TopBP1 ^12–14^, and POLQ ^39^ in controlling RAD51 loading to chromatin, and the role of RAD51 in DSB repair ^12, 13^, the *Cip2a*−/− mammary epithelial cells exposed to IR displayed enhanced RAD51 chromatin recruitment (Fig. 4I,J). The enhanced chromatin recruitment of both TopBP1 and RAD51 in CIP2A-deficient cells provides as mechanistic explanation for the observed G2/M cell cycle arrest ^8, 10, 11, 40, 41^ (Fig. 1G). Finally, it has been reported that TopBP1 determines cancer cell sensitivity to PARP inhibition by regulating RAD51 chromatin loading ^13^. Also consistent with this observation, *CIP2A* depletion hypersensitized BRCA-proficient MDA-MB-231 cells to two different PARP inhibitors (Fig. S4C).

**Figure 4:**
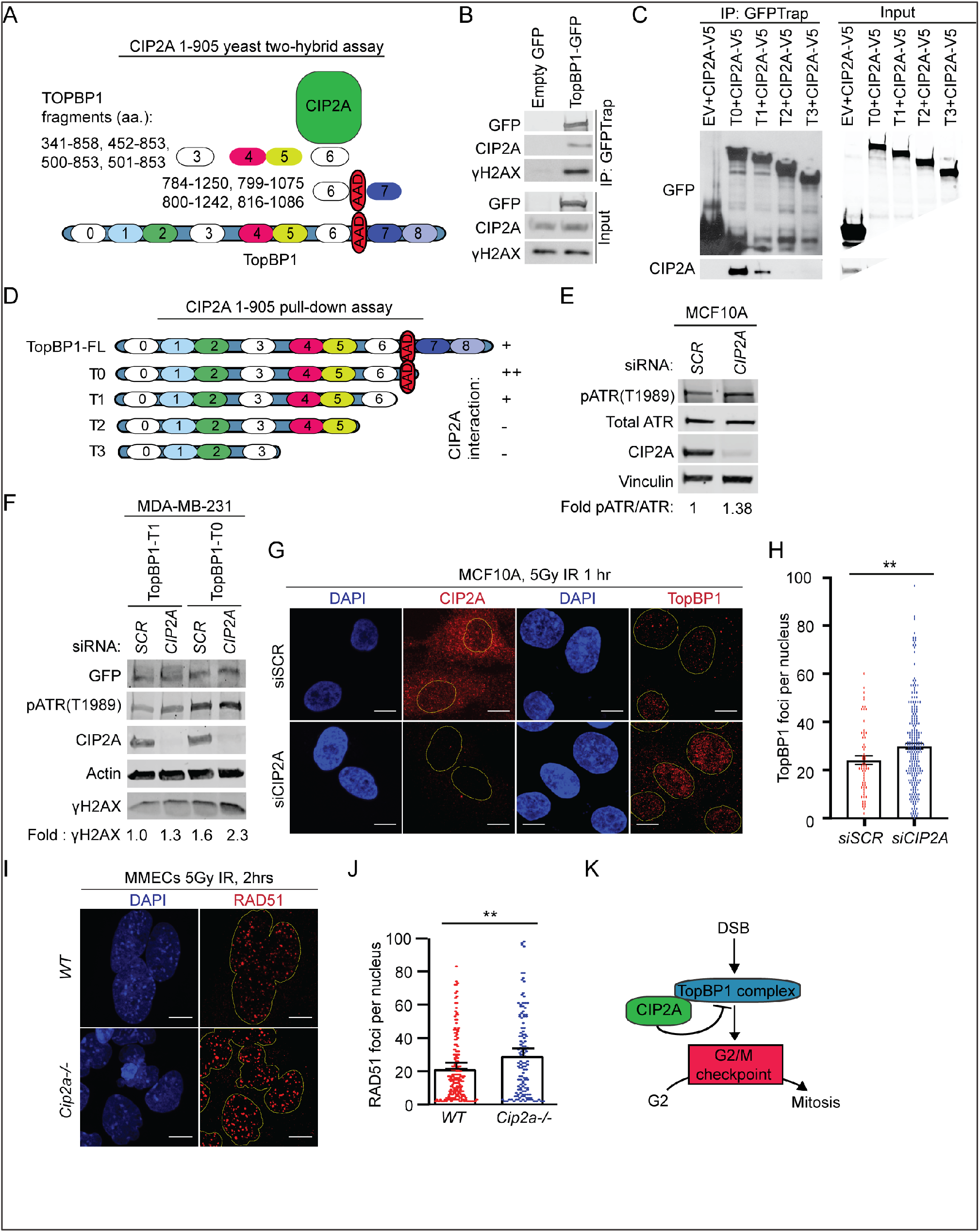
CIP2A is an interacting partner of TopBP1 and promotes mitotic progression of DNA damaged cells. **A**, Schematic presentation of breast cancer cell line cDNA fragments coding for TopBP1 domains that interact with full length CIP2A in a yeast two-hybrid assay. Numbers in the TopBP1 drawing refer to BRCT domains 1-8; AAD, ATR activation domain. **B**, Co-immunoprecipitation between endogenous CIP2A and ɣH2AX in HEK293 cells transiently overexpressing GFP or full length TopBP1-GFP as indicated. Input 5% of total IP. **C**, Co-immunoprecipitation of CIP2A in HEK293 cells transiently overexpressing V5-tagged CIP2A and GFP-tagged Empty vector (EV) or TopBP1 truncated mutants T0, T1, T2, T3 as indicated in (D). Input 5% of total IP. **D**, Schematic representation of TopBP1 mutants used in (B,C) Relative interaction efficiencies are estimated from the experiment where all indicated mutants were included. **E**, Basal-like immortalized MCF10A cells transfected with non-targeting (*SCR*) or *CIP2A* siRNAs for 48hrs. Immunoblot of whole cell extracts (WCEs) probed for pATR, total ATR and CIP2A. Vinculin was used as a loading control. Quantification represent mean of three experiments. **F**, MDA-MB-231 cells transfected with non-targeting (*SCR*) and *CIP2A* targeting siRNAs for 72 hrs and overexpressing TopBP1 mutants T0 and T1 as indicated for 48 hours. Immunoblot of WCEs probed for pATR, ɣH2AX and CIP2A. Actin was used as a loading control. Quantification represents mean of three experiments. **G**, IR-induced TopBP1 foci formation in MCF10A cells transfected with *SCR* or *CIP2A* siRNA as indicated for 48 hrs. Cells were treated with 5Gy radiation for 1 hour and stained for CIP2A or TopBP1. **H**, Quantifications of the nuclear foci from (G) expressed as mean ± SD from representative experiment of three experiments with similar results **I**, IR-induced RAD51 foci formation in mouse mammary epithelial cells (MMECs) isolated from *WT* and *Cip2a*−/− mice cultured *in-vitro* for 48 hrs, treated with 5Gy radiation for 2 hours. Images were taken at 63X on 3i spinning disk confocal and at least 150 cells quantified per each condition using speckle counter pipeline on Cell Profiler. Scale bar: 10μM. **J**, Quantifications of the foci in expressed as mean ± SD of representative experiment. All statistical analyses were conducted with Welch’s Student t-test for unequal variances, *p<0.05, ** p<0.01, ***p<0.001. **K**, Schematic presentation of the role of CIP2A in inhibiting TopBP1-elicited G2/M checkpoint activation.

Collectively, the newly discovered role for CIP2A in blunting TopBP1 and RAD51 chromatin recruitment provides a mechanism for dampening of the DDR ^5^, and G2/M checkpoint, in DNA-damaged cells (Fig. 4K). DDR dampening also provides a plausible mechanistic explanation for the requirement of CIP2A for continuous proliferation of DNA-damaged mammary epithelial cells; and thereby for BLBC initiation.

### Clinical and functional relevance for CIP2A in human BLBC

In concert with the other results, across the human breast cancer subtypes, *CIP2A* mRNA was found to be highest expressed in BLBC (Fig. 5A and S5A). Notably, also *TOPBP1* was highest expressed in BLBC subtype (Fig. S5B). Although regulation of *CIP2A* expression in BLBC has not been studied, overexpression in BLBC is most likely a result of the fact that EGFR expression is the determining hallmark of BLBCs ^1, 2^ and that EGFR in known to positively regulate *CIP2A* gene expression ^21, 22^. In addition, there is a very high prevalence of *TP53* mutations in BLBC which results in activation of *CIP2A* gene promoter activity through the p21-E2F1 pathway ^20^. Consistent with these observations, a significant correlation between *TP53* mutation, and high *CIP2A* expression was confirmed in the GSE21653 cohort (Fig. S5C). Furthermore, *CIP2A* gene promoter activity is known to be stimulated by DNA-PK ^42^, which is overactive in BRCA-deficient cells ^3^. The clinical relevance of CIP2A in BLBC was also evident from patient survival analysis. Both high mRNA and protein expression of CIP2A predicted poor disease-free or overall survival only in BL-TNBC, but not in non-BL-TNBC, or among unselected breast cancer patients (Fig. 5B-D, and S5D-H). Notably, the 5-year survival of patients with highly CIP2A positive BL-TNBC tumor was only about 50% in both patient cohorts (Fig. 5B,D), indicating that these tumors are particularly aggressive.

**Figure 5:**
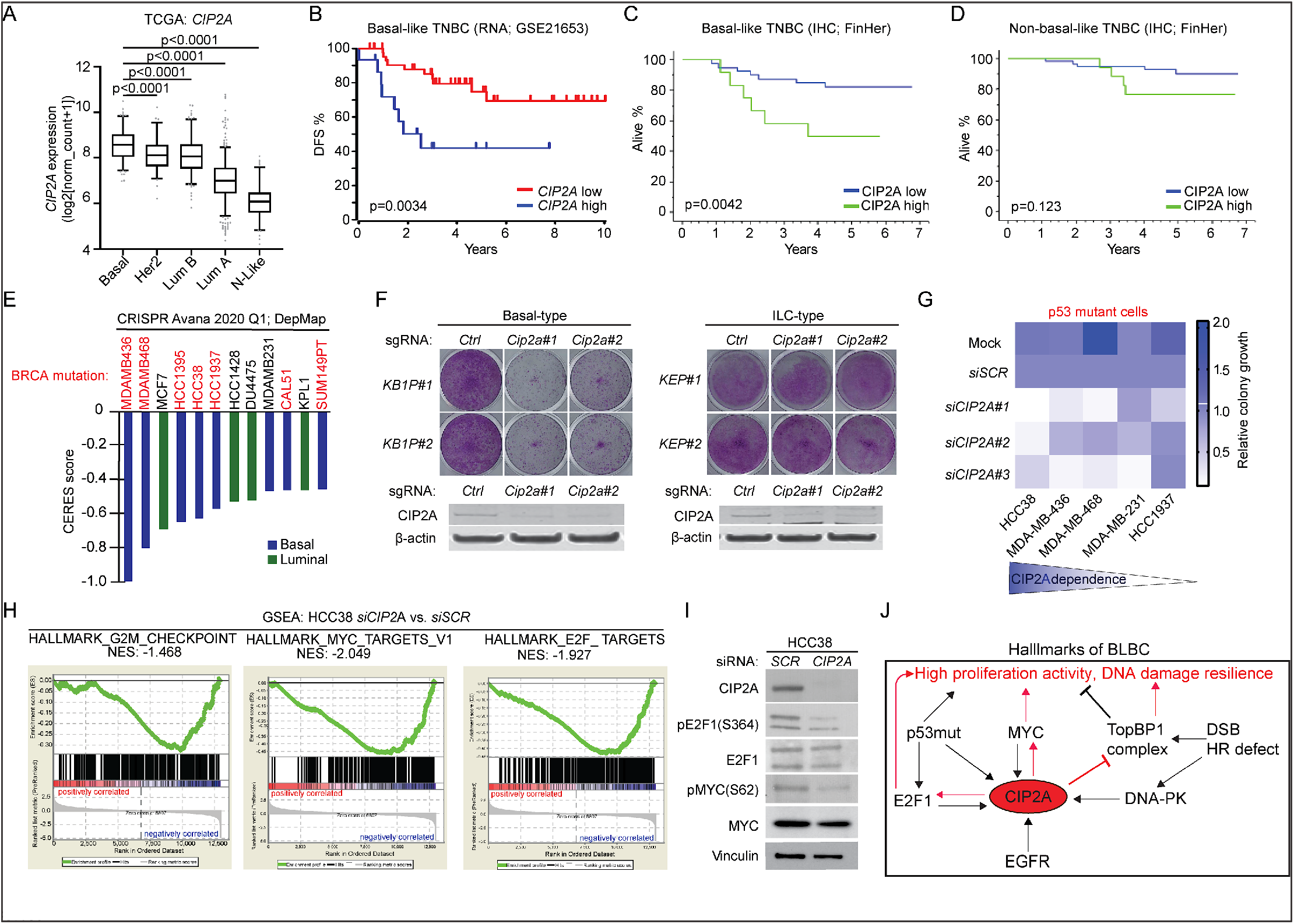
CIP2A associates with poor prognosis and drives growth of BLBC cells. **A**, Expression of *CIP2A* mRNA in indicated molecular subtypes. Data derived from TCGA. P-values by unpaired t-test. **B**, Disease-free survival of *CIP2A* high (n=15) and *CIP2A* low (n=45) expressing basal-like TNBC patients in GSE21653 cohort. **C**, Overall survival of CIP2A high (n=12) and CIP2A low (n=51) basal-like TNBC patients in FinHer cohort. **D**, Overall survival of CIP2A high (n=17) and CIP2A low (n=47) non-basal like TNBC patients in FinHer cohort. **B-D**, P-values calculated by log-rank test. **E**, *CIP2A* dependence of breast cancer cell lines with CERES score < −0.4 from DepMap portal (Avana 2020Q1). Lower CERES scores indicate that the cell line is more dependent on *CIP2A*. Color coding indicates the breast cancer subtype of the cell line based on PAM50 classification. **F**, Colony growth assays conducted on mammary tumor cell lines isolated from basal-type (*KB1P#1* and *KB1P#2*: *Brca1* and *Trp53* mutant) and invasive lobular carcinoma (ILC)-type (*KEP#1* and *KEP#2*: *E-Cadherin* and *Trp53* mutant) mouse models; *Cip2a* was knocked out using CRISPR/Cas9 using 2 unique gRNAs. Western blots from the same samples probed for CIP2A below. Shown are representative images of at least 2 independent biological repeats for each cell line. **G**, Summary of *CIP2A*-dependence on colony growth of indicated *TP53*-mutant TNBC cell lines transfected with Mock, non-targeting siRNA (*siSCR*), or three unique *CIP2A* targeting siRNAs (*siCIP2A #1, #2, #3*). Colony areas were quantified and normalized to *siSCR*. **H**, Gene Set Enrichment Analysis (GSEA) conducted on differentially expressed genes obtained from RNA-seq of HCC38 cells depleted with 3 unique *CIP2A* siRNAs. **I**, HCC38 cells transfected with *SCR* or *CIP2A* siRNAs for 72 hrs and immunoblotted for indicated protein. **J**, Schematic presentation of the role for CIP2A in coordinating BLBC molecular hallmarks. Red arrows indicate mechanisms by which CIP2A drives BLBC initiation and progression. Black arrows indicate known mechanisms implicated in BLBC that either increase CIP2A expression and/or promote BLBC progression.

To functionally assess BLBC cell dependence on *CIP2A*, we first surveyed the Dep-Map essentiality database across 33 breast cancer cell lines classified according to PAM50 classification either to luminal, basal or HER2-positive. Among the 12 cell lines with CERES gene dependency score less than −0.4 for *CIP2A* loss, the majority of cell lines were found to be BLBC cells (Fig. 5E, Table S2). Notably, all except one of these most *CIP2A*-dependent BLBC cells carried either a *BRCA1* or *BRCA2* mutation (Fig. 5E). To further substantiate these results, in a genetically defined CRISPR/Cas9 model, *Cip2a* was found to be essential for colony growth of mouse mammary tumor cells depleted for *Trp53* and *Brca1* (*KB1P*; basal-type)^43^(Fig. 5F). However, *Cip2a* was dispensable for growth of either *Trp53/E-cadherin* mutant mammary tumor cells (*KEP*; invasive lobular carcinoma-type)^44^(Fig. 5F), or cells from the mice with activated AKT and loss of E-cadherin in mammary tumor cells (*WEA*; invasive lobular carcinoma-type)^45^(Fig. S5I). Further, RNA-sequencing analysis from the most *CIP2A*-dependent and *TP53*-mutant BLBC cell line (Fig. 5G, S5J) HCC38 (*TP53 mutant/BRCA1 promoter methylation/BRCA2* mutant), revealed that CIP2A drove a gene expression program that was consistent with its role as a BLBC driver. Specifically, CIP2A drives expression of G2/M-associated genes, as well as MYC and E2F1-driven gene expression programs (Fig. 5H). The role of CIP2A in inhibiting the dephosphorylation of the activating phosphorylation sites in both MYC and E2F1 was confirmed by western blot analyses (Fig. 5I).

These data strengthen the evidence for, and confirm the selective CIP2A-dependence of BLBC cells harboring genomic instability and HR defects. Consistent with being regulated by the key pathways of BLBC, namely MYC, E2F1, EGFR and DDR, our data reveals CIP2A as a BLBC protein driver comprehensively coordinating the molecular disease hallmarks of this disease subtype (Fig. 5J).

### Transcriptional CIP2A targeting by SMAPs as potential BLBC therapy

Effective treatment of BLBCs represents a significant unmet medical need as a result of both intrinsic and acquired chemotherapy resistance, as well as a lack of therapeutically targetable driver alterations. To credential the role of CIP2A as a BLBC drug target, we tested whether a recently developed series of <u>S</u>mall <u>M</u>olecule <u>A</u>ctivators of <u>P</u>P2A (SMAPs) ^15, 16^, which have been shown to reactivate the CIP2A-targeted PP2A complex (PP2A-B56)^15, 46^, could be used to target CIP2A-expressing BLBC. SMAPs have thus far been shown to be effective against several MYC-driven cancer cell lines, including established TNBC cells ^47^, but there is no information whether their therapeutic action is related to CIP2A. To start with, we verified that treatment with two independent SMAPs (DBK-1154 and DT-061) resulted in a robust concentration-dependent inhibition of cell viability in eight established BL-TNBC cell lines (Fig. 6A and S6A). To ask whether SMAPs are effective against patient-derived cells, we used the recently characterized five BLBC patient-derived cancer stem cell-like lines ^48^. Notably, consistent with notion that these cell lines were derived from tumors of patients that had undergone neoadjuvant chemotherapy, all five cell lines showed resistance to classical chemotherapies (Fig. 6B). However, regardless of their chemoresistance, these CIP2A positive (Fig. S6B) patient cells retained their sensitivity against all three tested SMAPs (DBK-1154, DT-061, NZ-1160)(Fig. 6B). Further, we used an orthotopic PDX model from a patient with *TP53* mutant, EGFR+ BLBC that was propagated *in vivo* to validate the therapeutic potential of our findings. Upon establishment of tumors, the mice were orally treated with DT-061, and tumor growth was measured. Directly supportive of their therapeutic relevance, oral DT-061 therapy resulted in significant inhibition of PDX growth over the 40-day treatment period (Fig. 6C). Similar to other *in vivo* studies with SMAPs ^15, 47, 49, 50^, we did not observe any treatment-related adverse effects in mice. Importantly the control tumors were CIP2A positive whereas tumors from DT-061 treated mice showed a clear trend for reduced CIP2A protein levels (Fig. S6C,D). These results clearly indicate that pharmacological PP2A reactivation could represent a novel therapeutic strategy for the treatment of therapy-resistant and CIP2A positive BLBCs.

**Figure 6:**
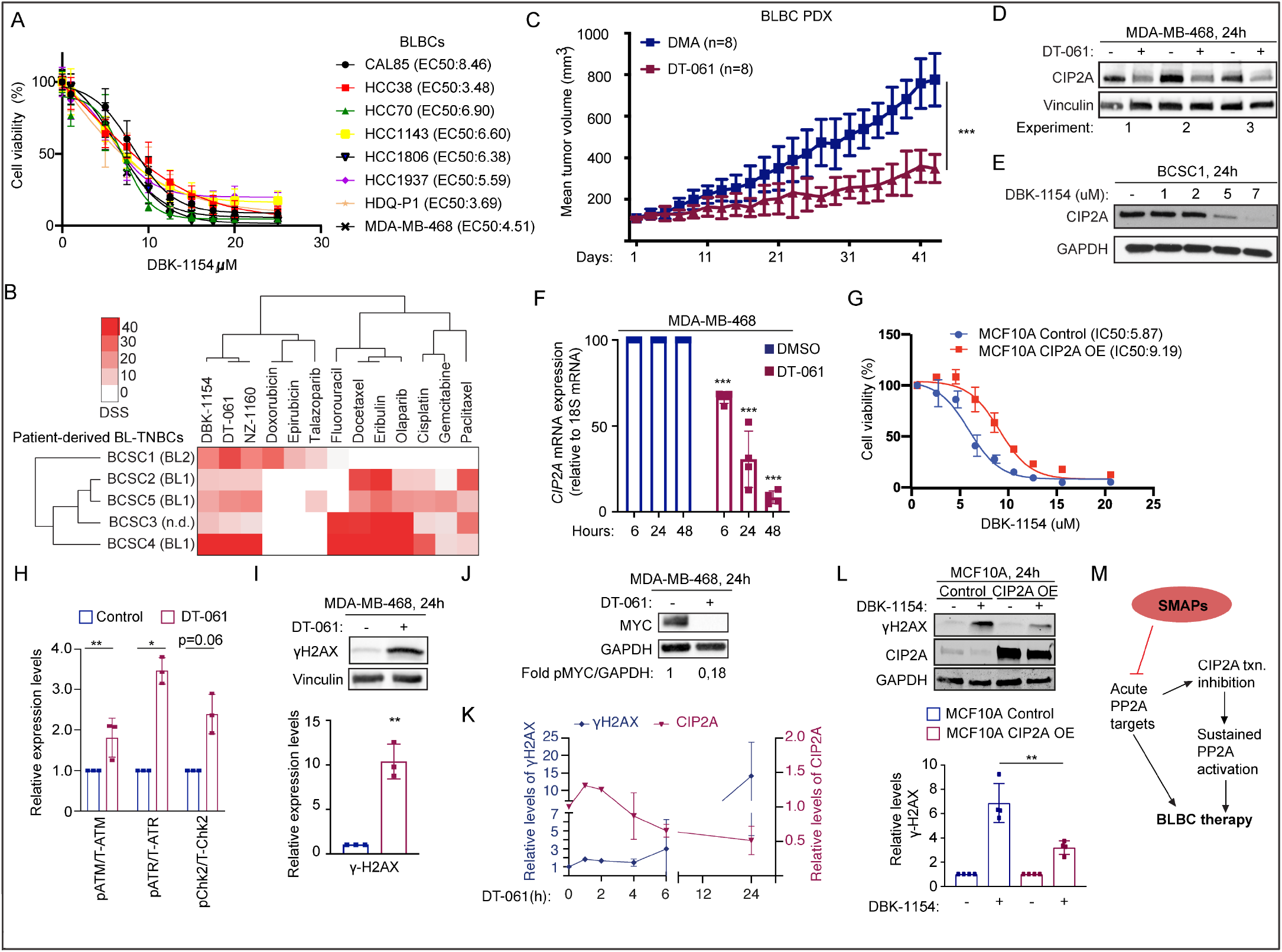
CIP2A targeting by SMAPs as potential BLBC therapy. **A**, SMAP (DBK-1154) sensitivity profiles of eight BL-TNBC cell lines. Cell viabilities were measured using CellTiterGlo Luminescence Assay after 24 hrs of drug treatment. EC50s are listed in parentheses **B**, Screening of patient-derived BLBC stem cell like cells for chemotherapy and SMAP responses. Heatmap indicates the drug sensitivity scores (DSS) of these cells across standard chemotherapeutics and three SMAPs DBK-1154, DT-061, NZ-1160). Higher DSS value indicates higher sensitivity. **C**, Tumor growth of an orthotopic patient derived xenograft model of basal triple negative breast cancer treated with DMA or 5mpk BID SMAP DT-061 for 43 days. Respective quantifications are represented as mean ± SD. **D,E**, SMAP treatment leads to CIP2A depletion. CIP2A western blots from MDA-MB-468 (D) and patient-derived stem cell-like cells (14-72)(E) on treatment with indicated SMAPs for 24h. DT-061 and DBK-1154 concentration 20 μM. **F**, Kinetics of *CIP2A* mRNA expression from MDA-MB-468 cells after treatment with 20uM of DT-061. n=3 expressed as mean ± SD. **G**, Dose response curve of control and CIP2A OE stable cell line (CIP2A OE) MCF10A cells on treatment with concentration series of DBK-1154 for 24 hours. IC50 values indicated in parentheses. **H**, Quantification of western blots displayed in S7A expressed as mean ± SD from n=3 replicates normalized to the untreated controls. **I,J**, Western blots of MDA-MB-468 cell line treated with 20μM SMAP DT-061 for 24 hrs and probed for ɣH2AX and MYC; ɣH2AX quantifications from n=3 replicates displayed below (I). Values for MYC represent mean of two experiments **K**, Time course of CIP2A and ɣH2AX protein expression in MDA-MB-468 treated with DT-061 (20 μM) for indicated time periods. Western blot data are shown in Fig. S7B. **L**, CIP2A overexpression in MCF10A cell line rescues the SMAP-elicited ɣH2AX activation effects. Western blots of parental and CIP2A OE MCF10A cells treated with SMAP DBK-1154 for 24hrs and probed for ɣH2AX. GAPDH is used as loading control; ɣΗ2AX quantifications from n=4 replicates displayed below. **A-L**, p-values calculated using unpaired t-test, *P<0.05, **P<0.01, *** P<0.001, ****P<0.0001. **M**, Schematic model of bi-phasic therapeutic action of SMAPs in BLBCs. SMAP treatment of cells results in acute inhibition of PP2A phosphotargets involved in both CIP2A regulation and BLBC growth. The acute SMAP response is sustained by PP2A reactivation resulting from SMAP-elicited CIP2A inhibition. txn; transcription.

Surprisingly, but related to potential link between SMAP response and CIP2A, western blot analyses revealed a potent inhibition of CIP2A protein expression by SMAPs at 24 hours in both the established cells, and in patient-derived cells (Fig. 6D,E and S6E-G). Indicative of transcriptional level regulation, CIP2A protein inhibition was accompanied with inhibition of *CIP2A* mRNA expression (Fig. 6F and S6G). Furthermore, rescue of CIP2A by exogenous overexpression shifted the SMAP IC50 response of basal-like immortalized MCF-10A cells (Fig. 6G, and S6H). SMAPs, albeit not equaling direct CIP2A inhibition, but representing now surrogate CIP2A inhibitors, were next tested for possible effects on biomarkers of CIP2A activity. Consistent with results in CIP2A-inhibited cells (Fig. 4), SMAPs induced potent checkpoint signaling exemplified by phosphorylation of ATR and H2AX, as well as phosphorylation of ATM and CHK2 (Fig. 6H,I and S7A). SMAP treatment also resulted in inhibition of MYC expression (Fig. 6J). Notably, especially p-ATR and γH2AX induction by SMAP occurred after CIP2A protein inhibition (Fig. 6K and S7D,E). Furthermore, CIP2A overexpression significantly rescued SMAP-elicited γH2AX induction (Fig. 6L). These results reveal that SMAPs have bi-phasic therapeutic activity consisting of direct PP2A activation ^15, 16^, followed by transcriptional inhibition of *CIP2A* expression discovered here (Fig. 6M).

## Discussion

Breast cancers are a heterogeneous group of malignant diseases. Whereas driver mechanisms and therapeutic strategies for steroid hormone receptor-positive cancers and HER2-positive breast cancers are more established; BLBCs, lack identified genetic drivers, and their therapies are often limited to relatively untargeted systemic therapies, such as conventional chemotherapy ^1, 2^. The lack of defined driver mechanism(s) is thus one important reason for overall poor patient survival especially in BLBCs. In this study we provide compelling cell based and *in vivo* evidence for a central role for CIP2A as a non-genetic driver of BLBC initiation and progression, and identify SMAPs as potential novel therapy for aggressive CIP2A positive BLBC tumors.

*CIP2A* gene sequence is not altered in BLBCs (https://cancer.sanger.ac.uk/cosmic). Instead, its expression is enhanced due to constitutive DDR activity ^42, 51^, TP53 inactivation ^20^, and EGFR pathway activation ^21^, which are all molecular hallmarks of BLBC^1, 3^. Our data expand on these findings by demonstrating induction of *Cip2a* mRNA expression in premalignant mammary tissue of DMBA-treated mice (Fig. 2A). This can be explained by the aforementioned DDR activity, but also by DMBA-induced activation of other pro-tumorigenic pathways such as the MEK-ERK pathway and MYC (Fig. 2B), known to stimulate *CIP2A* transcription ^18^. Therefore, transcriptional *CIP2A* induction early in DMBA-induced tumorigenesis is fully supportive of its role as a BLBC driver essential for tumor initiation. Later in the human BLBC progression when TP53 is lost, *CIP2A* transcription is permanently enhanced by increased p21-E2F1 activity ^20^. Together these findings provide an explanation for high *CIP2A* expression in BLBC (Fig. 5A), and a functional link between two human major tumor suppressors, TP53 and PP2A.

Although PP2A inhibitor proteins have oncogenic functions, none of them, including CIP2A, have been shown to be essential for *in vivo* tumorigenesis. As opposed to previous assumptions that CIP2A is involved in the development of multiple human solid cancers ^18^, including breast cancers, our results demonstrate striking selectivity in essentiality for CIP2A in the initiation and progression of BLBCs. Whereas *Cip2a* was required for DMBA-induced mouse BLBC initiation, it was not essential for initiation of DMBA-induced lung, ovary, skin, or stomach tumors (Fig 1D). We further confirmed that *Cip2a* was dispensable for skin and ovarian tumorigenesis by independent *in vivo* models. Further, in genetically defined cell culture models, the *Brca1/Trp53* mutant basal-like cells, but not the invasive lobular carcinoma-type mouse mammary tumor cells were dependent on *Cip2a* for their colony growth. Of clinical relevance, in human breast cancer samples, both at mRNA and protein level, high CIP2A expression predicted for poor patient survival exclusively in BLBCs, but not in other studied breast cancer subtypes. Importantly, our results also provide a plausible mechanistic explanation for the noted dependency of CIP2A both for initiation and progression in BLBCs. We note that *CIP2A* is not only itself regulated by BLBC hallmarks, but also controls many of the molecular hallmarks of BLBC^1, 3^, including survival promotion of *TP53/BRCA*-deficient cells, high MYC and E2F1 transcriptional activity, as well as resilience of cell proliferation under a high degree of DNA damage (Fig. 5J). Although these are also important mechanisms for HGSOC development, our results suggest that CIP2A has differential effects on DDR proteins essential for BLBC or HGSOC. Together these data provide compelling evidence to support discovery of CIP2A as a specific BLBC driver protein implicated both in tumor initiation and progression.

Through genome-wide dependence mapping, CIP2A was identified as a functional homologue for several critical DNA damage proteins. Although it is very likely that also the other DDR proteins identified in the functional network with CIP2A contribute to phenotypes in CIP2A-deficient cells, we focused on validation of functional interaction between CIP2A and TopBP1. We demonstrate both a direct protein interaction between CIP2A and TopBP1 and that CIP2A prevents retention of TopBP1 and RAD51 on damaged chromatin in premaligant basal-like mammary cells. Thus, in *CIP2A* deficient cells TopBP1 can induce effective DDR, whereas in pre-malignant *CIP2A* positive cells the DDR is dampened which allows for continued mitotic activity (Fig. 4K)^8, 10, 11, 40, 41^. Importantly the link between CIP2A and DDR may also provide a plausible explanation for the dilemma that PP2A inhibition should not be essential for tumorigenesis in mouse cells ^52^. While it has been convincingly shown that PP2A inhibition is not required for mouse cell transformation by hyperactivated RAS ^52^, CIP2A’s role in DNA-damage-induced cell transformation may not follow the rules of RAS-dependent transformation. Specifically, even though CIP2A-mediated PP2A inhibition supports phosphorylation of MYC, E2F1 and NBN relevant to this study ^20, 35, 53^, it is possible that CIP2A-mediated BLBC initiation is not fully dependent on PP2A inhibition, but may also result from CIP2A’s role as a direct TopBP1 interacting protein, and consequent PP2A-independent effects of CIP2A on TopBP1 function.

In addition to identifying CIP2A as a protein driver for BLBC, we demonstrate that a first-in-class series of small molecule activators of PP2A (SMAPs)^15, 16^, function as inhibitors of *CIP2A* expression. Our results reveal a model where SMAPs initially directly activate PP2A-B56 ^15^, and lead to a prolonged response by transcriptional downregulation of the PP2A-B56 inhibitor *CIP2A* (Fig. 6M). The mechanisms for SMAP-elicited *CIP2A* mRNA inhibition has yet to be elucidated, but as *CIP2A* promoter activity is stimulated both by MEK-ERK-ETS pathway ^21^, and by MYC ^54^, and SMAPs inhibit both ERK activity and MYC (Fig. 6J), these findings provide a plausible mechanistic explanation for the *CIP2A* mRNA inhibition by SMAPs. Importantly, we were also able to demonstrate that CIP2A overexpression partly rescued the effects of SMAPs as assessed by both cell viability and DDR regulation. However, it is important to note that we consider SMAPs as surrogate CIP2A inhibitors that also have acute effects not mediated by CIP2A inhibition ^15^, thereby explaining the effects of SMAPs also in other types of cancer cells ^47, 49, 50^. Importantly, we validated the therapeutic effect of several SMAPs across 15 different cell lines, including 6 individual patient-derived lines and a PDX model, together minimizing cautions related to known intratumoral heterogeneity of BLBC tumors ^55^. The effects on MCF-10A cells also indicate potential usefulness of SMAPs in eradicating the low transformation level basal-like mammary epithelia cells. Our xenograft data provide first evidence for *in vivo* efficacy of SMAPs on patient-derived BLBC cells. However, this is directly supported by recent data demonstrating that both CIP2A inhibition, and SMAP treatment can significantly inhibit xenograft growth of established TNBC cell lines ^28, 32, 47^. Based on our results with PARP inhibitors, and recent studies implicating that PP2A reactivation potentiates the therapeutic effects of numerous different types of drugs ^35, 50^, future studies should be directed towards comprehensive screening efforts to the most efficient combination therapies with SMAPs for patients with aggressive CIP2A positive BLBC tumors. Another very interesting future direction would be to test the effects of SMAPs on brain metastasis of BLBCs, as SMAPs were recently shown to cross blood-brain-barrier, and to induce significant survival effects in an intracranial glioblastoma model ^49^.

Together these results credential a therapeutically actionable driver protein for one of the most aggressive human cancer types, BLBCs. We also discover novel link between CIP2A and DDR via direct interaction with TopBP1. More generally, these results emphasize the importance in characterizing and functionally validating protein level dysregulation of key signaling effectors in cancer types for which apparent genetic drivers are lacking.

## Materials and Methods

### Mouse experiments

In order to develop DMBA-induced tumors in *WT* and *Cip2a*−/− female mice, they were administered with 1mg of DMBA dissolved in 200μl of corn oil by oral gavage once a week for 6 weeks starting at 12-14 weeks of age as previously described ^26^. The mice were monitored twice a week for tumor formation until morbidity. Mice were sacrificed upon tumor burden and/or when they showed general signs of illness. Upon autopsy tumors in different tissues were recorded and collected. To analyze DMBA-induced mutation load and *Cip2a* mRNA expression in *WT* and *Cip2a*−/− premalignant mammary gland tissues, the mice were sacrificed 2 weeks after the last DMBA treatment and tissues were collected for further analysis. DMBA/TPA protocol for skin tumorigenesis and experiments with *Cip2a+/−* mice crossed with an ovarian cancer mouse model *TgMISIIR-Tag* are described in supplementary materials and methods. Tissue samples collected for extraction of RNA and genomic DNA were snap frozen into liquid nitrogen. Tissue samples for histochemical and for immunohistochemical analysis were fixed in formalin.

Mouse mammary epithelial cells (MMECs) were isolated from 3 to 4 months old *Cip2a*−/− and *WT* mice and cultured *in vitro* as described in ^56^. Briefly, mammary glands (without lymph nodes) from 3-4 mice per genotype were pooled together in cold PBS, minced using scalpels and collected to warm collagenase solution. The samples were agitated for 2 to 3 hours at 37°C and resuspended in DMEM/F12 isolation medium containing 20 U/ml DNAse I. They were subjected to a few rounds of pulse centrifugations (1500G) until contaminating red blood cells (RBCs) disappeared from the pellet. The final clear cell pellets (containing mammary epithelial ducts) were dissociated into single cells using Accutase (StemCell Technologies). The obtained single cells were cultured using DMEM/F12 culture medium for IF experiments or used directly for flow cytometry. The recipes of the different media used in the process are listed and described in Table S3.

Mouse tumor cell lines were generated from spontaneous mammary tumors of following breast cancer mouse models: *K14Cre; Brca1^F/F^; Trp53^F/F^*(*KB1P*) ^43^, *K14Cre; Cdh1^F/F^; Trp53^F/F^*(*KEP*) ^44^ and *Wap-cre; Cdh1^F/F^; Akt1^E17K^*(*WEA*) ^45^. Tumor cell lines were generated by collecting tumors in cold PBS and minced by chopping with scalpels. Aggregates were plated out. *KEP* and *WEA* tumor cell line cultures were incubated at 37°C with 5% CO_2_ and 20% O_2_. *KB1P* cell lines were incubated at 37°C with 5% CO_2_ and 3% O_2_. Homogenous epithelial cell morphology was obtained after cultures were passaged 2-3 times. Used cell culture media are described in Table S3.

### Cell culture and transfections

All the commercial cell lines used in this paper were purchased from American Type Culture Collection (ATCC) or Leibniz Institute’s German Collection of Microorganisms and Cell Cultures (DSMZ). All the cells in culture were negative on periodically testing for mycoplasma using Mycoplasma Detection Kit (Roche). All the human and mouse cells, their culture conditions and supplements used for cell culture are listed in Table S3. Breast cancer stem-like cells (BCSCs) were isolated from TNBC patients who received standard chemotherapy and cultured as described previously ^48^. MCF10A stable cell lines overexpressing CIP2A-V5 and empty vector (MCF10A-CIP2A OE and MCF10A-Control) were generated using the lentiviral constructs pWPI-CIP2A-V5 and pWPI respectively. After transduction with lentiviral particles, successfully transfected (GFP positive) cells were sorted using SH800 Cell Sorter (Sony). Plasmid DNAs and siRNAs were transfected using Jet Prime (Polyplus Transfection) and Oligofectamine (Thermo Fisher Scientific) reagents respectively as per manufacturer’s protocols. DNAs were transfected for 48 hours and siRNAs were transfected for 48 to 72 hours until use for further experiments.

### CRISPR/Cas9 mediated gene disruption

In order to knockout *Cip2a*, mouse mammary tumor cell lines were transduced with lentiviral vectors lentiCas9-Blast carrying Cas9 and with lentiGuide-Puro containing sgRNA against mouse *Cip2a* or a control non-targeting (NT) sgRNA. Plasmid details in Table S4. The used two sgRNA sequences against *Cip2a* were selected from a genome-wide library of guideRNAs (Genome-scale CRISPR Knock-Out (GeCKO) v2.0)^57^. Cloning of sgRNAs into lentiGuide-Puro vector was performed as previously described ^58^. Cloned vectors were verified by Sanger sequencing. After selection of lentiCas9-Blast transduced cells, they were transduced with and selected for lentiGuide-Puro. Knockout efficiency was determined by analyzing CIP2A protein expression by western blot.

### Antibodies, RNAs, primers, and DNA constructs

Antibodies (along with dilutions for each application), plasmids and sequences of siRNAs, gRNAs and primers used are listed in Table S4.

### Co-immunoprecipitations

Co-IP experiments were conducted using the optimized protocols previously published for GFP tagged chromatin bound proteins, kindly provided by Prof. Andrew Blackford, ^59, 60^. Briefly, HEK293 cells were transfected with indicated plasmids for 48 hours. Cells were lysed using IP lysis buffer (containing 100mM NaCl, 1mM MgCl_2_, 10% glycerol, 0.2% Igepal CA630, 5mM NaF and 50mM Tris, pH 7.5) supplemented with 1X EDTA-free protease inhibitor tablet (Roche) and 25 units/ml Benzonase (Millipore) and rotated on a roller at 4°C for 20 minutes. After digestion of DNA and nuclear components, final concentration of NaCl and EDTA in the samples was adjusted to 200mM and 2mM respectively and rotated for another 10 minutes. The lysates were then cleared by high speed centrifugation (16000 rpm) for 15 minutes and 5% of the supernatant was kept aside as Inputs. The rest of the lysate was added to 20μl of GFP-Trap agarose beads (ChromoTek) and rotated on a roller at 4°C for 2-3 hours. The GFP bound complexes were washed 3-4 times and eluted using 2X Sample buffer. Protein interactions were assessed by western blot of Input and Co-IP samples.

### Immunofluorescence

MMECs and MCF10A cells were cultured in ibidi 8 well μ slides (ibiTreat #80826) for 24 hours. Cells were irradiated with 5Gy ionizing X-ray radiation (IR) using Faxitron Multirad 350. After the indicated time points, cells were fixed with 4% PFA for 15 minutes at room temperature (RT), permeabilized with 1%TritonX-100 in PBS for 15 minutes and blocked with 10% goat serum in PBS for 30 minutes. Primary antibodies were incubated overnight at 4°C and next day, Alexa Fluor conjugated secondary antibodies were incubated for 1 hour at room temperature. The Nuclei were counter stained using DAPI (Invitrogen). The nuclear foci were imaged using Zeiss LSM780 or 3i CSU-W1 Spinning disc confocal microscope (63X objective). Z-stack images were taken and maximum Z intensity projection images were used for image analysis. Nuclear foci were quantified using Speckle counter pipeline in Cell Profiler software ^61^. A minimum of 100 nuclei were counted for each condition. Each experiment was repeated with identical conditions 3 times.

### Mitotic index experiments

Mitotic index experiments were conducted by modifying previously published protocol described in ^8^. Briefly, MCF10A cells were transfected with indicated siRNAs for 24 hours, following which they were seeded into ibidi 8 well μ slides (ibiTreat #80826) for 24 hours. Cells were irradiated with 10Gy radiation followed by Nocodazole block (100ng/ml), one hour after IR for 18 hours. After the indicated time points, cells were stained for phospho-Histone H3 (Ser10) using similar immunofluorescence protocols as mentioned above. Images were taken on Zeiss Axiovert or EVOS fl Microscope with 10X objective and quantified using ImageJ software. Experiment was repeated 3 times.

### Protein isolation and western blotting

Protein lysates were prepared from cells by using RIPA buffer (50 mM Tris-HCl, pH 7.4, 150 mM NaCl, 1% NP-40, 0.5% DOC, 0.1% SDS, 2 mM EDTA) supplemented with protease and phosphatase inhibitors (Roche). Protein concentration was quantified using the BCA protein assay kit (Pierce). Equal amount of protein lysate was loaded with NuPage LDS Sample Buffer (ThermoFisher) onto 4–20% Mini-PROTEAN® TGX™ Precast Protein Gels (BioRad) or NuPage 4-12% Bis-Tris gradient gels (Invitrogen) and transferred onto Trans-Blot Turbo Midi Nitrocellulose membranes (BioRad) using Trans-Blot Turbo Transfer System (BioRad). Membranes were blocked with 5% milk in TBS-T or 10% Western Blot Blocking Reagent (Roche) followed by primary antibody incubation overnight at 4°C. Secondary antibodies were incubated for 1 to 2 hours at room temperature and membranes were imaged. For HRP antibodies detection was done using ECL based Curix 60 film processor (Agfa) and for IRDye Conjugated secondary antibodies Odyssey CLx imaging system was used.

### Colony formation assay

Optimized number of untransfected (Mock), non-targeting siRNA (*siSCR*) and 3 unique *CIP2A* targeting siRNA (*siCIP2A #1, siCIP2A#2, siCIP2A#3*) transfected HCC38, MDA-MB-436, MDA-MB-468, MDA-MB-231 and HCC1937 cells (2000-10,000 cells per well) were seeded in 12-well plates. In MDA-MB-231 cells, for testing PARP inhibitor sensitivity after *CIP2A* depletion, optimized number of *siSCR* and *siCIP2A* transfected cells were treated with indicated concentrations of PARP inhibitors (Olaparib and Niraparib) for 48 hours. After 7-10 days, colonies were fixed with cold methanol. Optimized number of control and *Cip2a* knock out *KB1P*, *KEP* or *WEA* cells (5000-20000 cells), were seeded into 12-well plates after transduction of and selection for lentiCas9-Blast and lentiGuide-Puro vector. After 5-7 days cell colonies were fixed and stained with 0.2% crystal violet solution prepared in 10% ethanol for 15 minutes at room temperature. Excess stain was removed by repeated washing with PBS or water. The colony areas were quantified using ColonyArea plugin ^62^ in Image J.

## Acknowledgements

We thank all Westermarck lab members for support during the project and especially Taina Kalevo-Mattila for technical assistance. Erica Nyman is thanked for help in IHC analysis. Professor Wojciech Niedzwiedz is thanked for GFP tagged-TopBP1 mutants and Dr. Andrew Blackford for sharing protocols. We are very grateful to Ruth Keri Lab from Case Western Reserve University for sharing the PDX model. Professor Johanna Ivaska is acknowledged for her valuable comments to the manuscript. We acknowledge important contributions by Finnish Functional Genomics Center, Cell Imaging and Cytometry core facility, and Genome Editing Core at Turku Bioscience Centre, and Turku Centre for Disease Modeling (TCDM) all supported by Biocenter Finland, and/or ELIXIR Finland. The Central Laboratory Animal Facilities of University of Turku are acknowledged for the genotyping, husbandry and exportation of the mice models used in this study. The project was funded by Academy of Finland (323096 for EP and 296801, 314443 and 310561 for LE), Finnish Cancer Foundations (JW), Finnish Cultural Foundation, Sigrid Juselius Foundation (JW, LE), Breast Cancer Now (JW, KW), and Governmental Research Funding for Turku University Hospital (TG). DCC is supported by the Fox Chase Cancer Center FCCC Core Grant NCI P30 CA006927. GN is supported by R01 CA181654, HL144741, CA240993, W81XWH-19-BCRP-BTA12 DOD and Rogel Cancer Gift Funds. JM is supported by Deutsche Forschungsgemeinschaft (DFG, German Research Foundation; PN(407869199)). AL was supported by Oncode Institute, Svenska Kulturfonden, Orion Research Foundation, Relander Foundation, Inkeri and Mauri Vänskä’s Foundation, Finnish Cultural Foundation’s Varsinais-Suomi Regional Fund and by K. Albin Johanssons Stiftelse.

