## Supplemental data for "CIP2A interacts with TopBP1 and is selectively essential for DNA damage-induced basal-like breast cancer tumorigenesis"

### CIP2A interacts with TopBP1 and is a protein driver for basal-like triple negative breast cancer

**This PDF file includes:**

Figures S1 to S7

Legends for Figures S1 to S7

Supplementary Materials and Methods

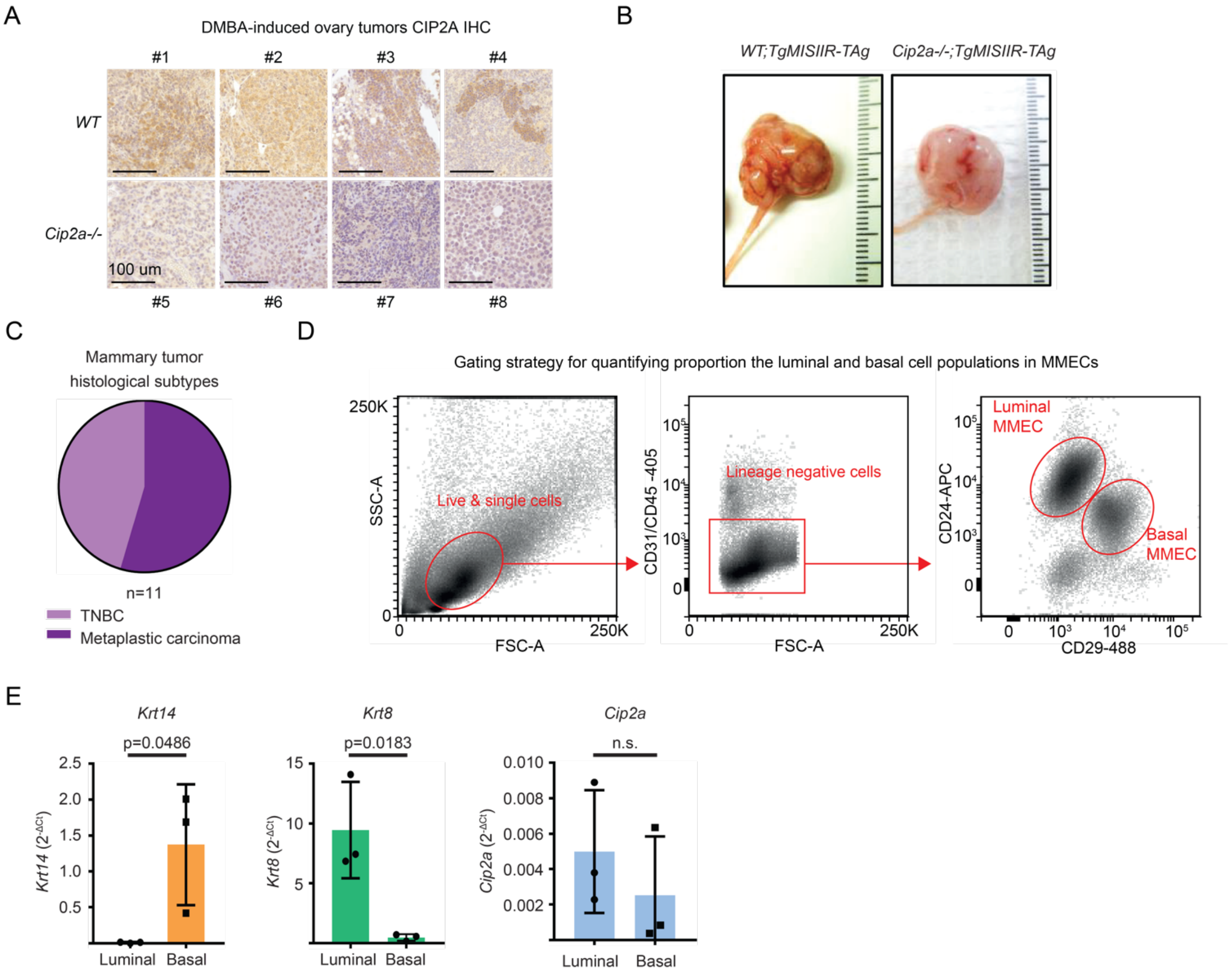


Figure S1

**Figure S1: A,** Representative images of immunohistochemical staining of CIP2A in DMBA-induced ovarian tumors in *WT* and *Cip2a-/-* mice. Scale bar: 100µm. **B,** Representative images of ovarian tumors collected from *WT;TgMISIIR-TAg* and *Cip2a-/-;TgMISIIR-TAg* mice. **C,** Histological classification of DMBA-induced mammary tumors from *WT* mice analyzed by a pathologist. **D,** Gating strategy to identify basal and luminal epithelial cell populations from mouse mammary gland tissue by flow cytometry. **E,** Analysis of basal marker *Keratin14* (*Krt14*), luminal marker *Keratin8* (*Krt8*) and *Cip2a* expression by qRT-PCR from flow cytometry sorted luminal and basal epithelial cell populations isolated from *WT* mouse mammary glands. *Krt14*, *Krt8* and *Cip2a* expressions normalized to *Gapdh*. p-values calculated by unpaired t-test.

A

Top25 *CIP2A* co-expressed genes (CCLE; 1156 cell lines)

| Rank | Correlated Gene | Cytoband | Spearman's Correlation | p-Value | q-Value |
|---|---|---|---|---|---|
| 1 | *SMC4* | 3q25.33 | 0.716 | 5.36E-182 | 1.70E-177 |
| 2 | *NCAPG* | 4p15.31 | 0.704 | 2.17E-173 | 3.43E-169 |
| 3 | *SGO2* | 2q33.1 | 0.675 | 2.14E-154 | 2.25E-150 |
| 4 | *CLSPN* | 1p34.3 | 0.668 | 4.04E-150 | 3.20E-146 |
| 5 | *SMC2* | 9q31.1 | 0.659 | 4.55E-145 | 2.88E-141 |
| 6 | *CENPE* | 4q24 | 0.654 | 6.53E-142 | 3.44E-138 |
| 7 | *CENPO* | 2p23.3 | 0.649 | 2.23E-139 | 1.01E-135 |
| 8 | ***POLQ*** | 3q13.33 | 0.646 | 2.26E-137 | 8.95E-134 |
| 9 | *KIF11* | 10q23.33 | 0.645 | 5.01E-137 | 1.76E-133 |
| 10 | *KIF14* | 1q32.1 | 0.636 | 3.80E-132 | 1.20E-128 |
| 11 | *BUB1* | 2q13 | 0.630 | 1.23E-128 | 3.53E-125 |
| 12 | *ARHGAP11A* | 15q13.3 | 0.629 | 2.23E-128 | 5.89E-125 |
| 13 | *BUB1B* | 15q15.1 | 0.628 | 6.09E-128 | 1.48E-124 |
| 14 | *PLK4* | 4q28.1 | 0.626 | 6.76E-127 | 1.53E-123 |
| 15 | *ZWILCH* | 15q22.31 | 0.624 | 6.70E-126 | 1.41E-122 |
| 16 | *GTSE1* | 22q13.31 | 0.621 | 2.81E-124 | 5.56E-121 |
| 17 | *KIF20B* | 10q23.31 | 0.618 | 9.42E-123 | 1.75E-119 |
| 18 | *KNL1* | 15q15.1 | 0.617 | 4.88E-122 | 8.57E-119 |
| 19 | *FANCB* | Xp22.2 | 0.616 | 1.72E-121 | 2.75E-118 |
| 20 | *ERCC6L* | Xq13.1 | 0.616 | 1.74E-121 | 2.75E-118 |
| 21 | ***TOPBP1*** | 3q22.1 | 0.615 | 1.95E-121 | 2.95E-118 |
| 22 | *CENPI* | Xq22.1 | 0.615 | 3.05E-121 | 4.39E-118 |
| 23 | *DLGAP5* | 14q22.3 | 0.614 | 5.80E-121 | 7.99E-118 |
| 24 | *CCNA2* | 4q27 | 0.614 | 7.22E-121 | 9.53E-118 |
| 25 | *ASPM* | 1q31.3 | 0.610 | 1.07E-118 | 1.36E-115 |

B

Basal-like breast cancers (n=142)

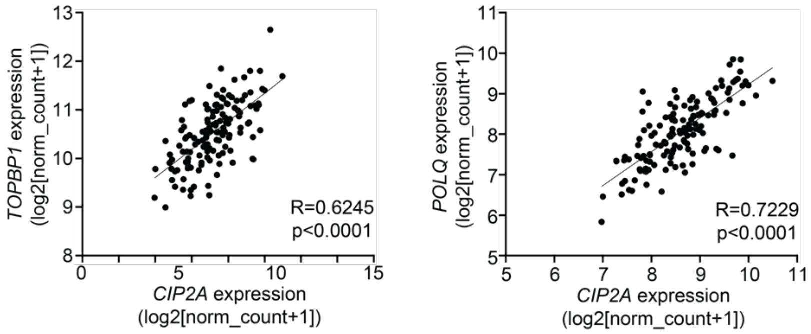


Figure S2

**Figure S2: A,** Top25 *CIP2A* co-expressed genes from mRNA co-expression analysis across 1156 samples with mRNA data from Broad Institute's Cancer Cell Line Encyclopedia (CCLE 2019). Data extracted from cBioportal (https://www.cbioportal.org/). **B,** Correlation of *CIP2A* and *TOPBP1* and *CIP2A* and *POLQ* gene expressions in basal-like breast cancer (n=142). Data derived from TCGA. R- and p-values calculated by Spearman correlation test.

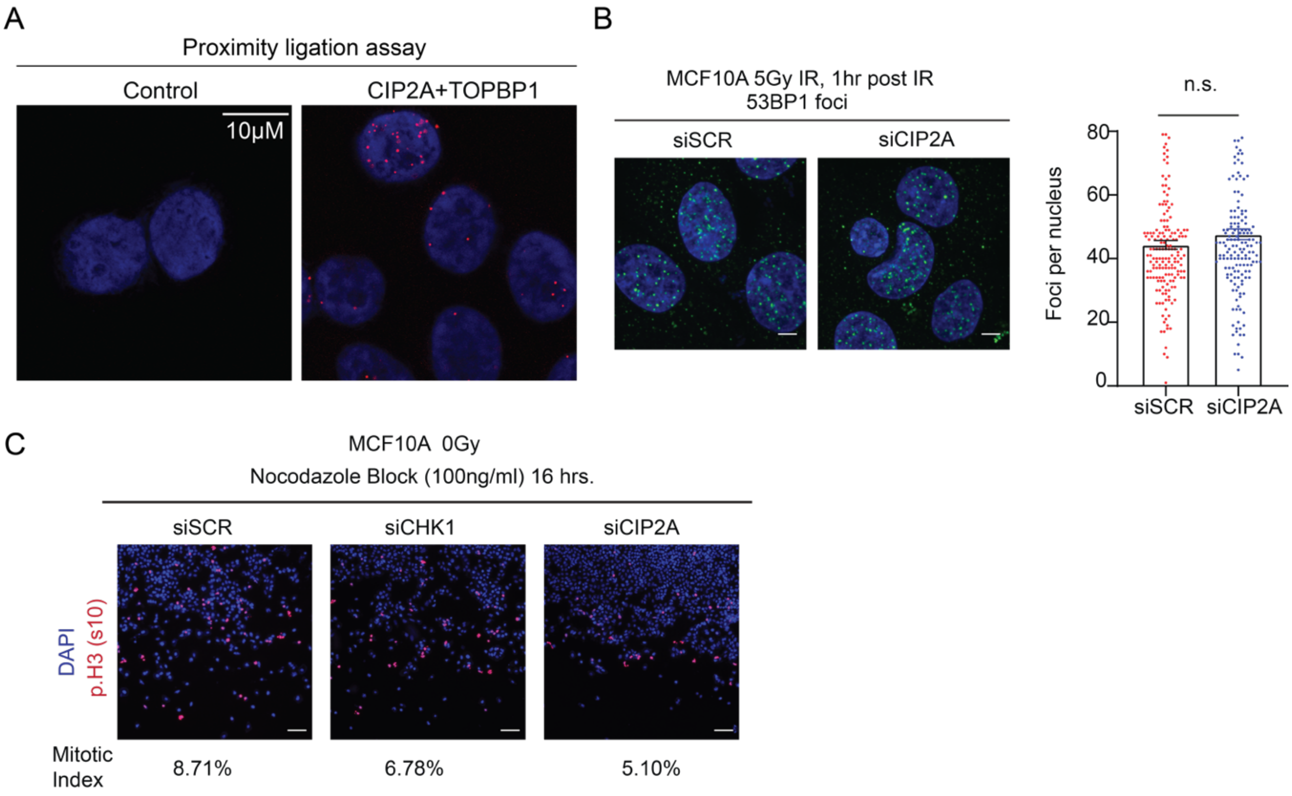


### Figure S3

**Figure S3: A,** Proximity Ligation assay (PLA) in HeLa cells in samples without primary antibodies (Control) and CIP2A Rabbit and TopBP1 mouse primary antibodies (CIP2A+TOPBP1). Red dots indicate proximity of CIP2A and TopBP1 proteins in the samples. Scale bar: 10μM. Imaging done using 63X magnification on Zeiss LSM780 confocal microscope. **B,** IR-induced 53BP1 foci formation in MCF10A cells transfected with *SCR* or *CIP2A* siRNA as indicated for 48 hrs. Cells were treated with 5Gy radiation for 1 hour and stained for 53BP1. Quantifications of the nuclear foci expressed as mean ± SD from representative experiment of three experiments with similar results. p-value calculated using unpaired t-test. **C,** Mitotic index analysis of MCF10A cells transfected with the indicated siRNAs. *CHK1* siRNA was used as a positive control. Cells were treated with Nocodazole (100 ng/ml) block without any IR for 16 hours. Mitotic cells were stained using phospho-histone H3 at Ser10. DAPI was used as counter stain. Scale bar: 100μM. Mitotic index (% of p.H3 Ser10 positive cells) indicated below the representative images.

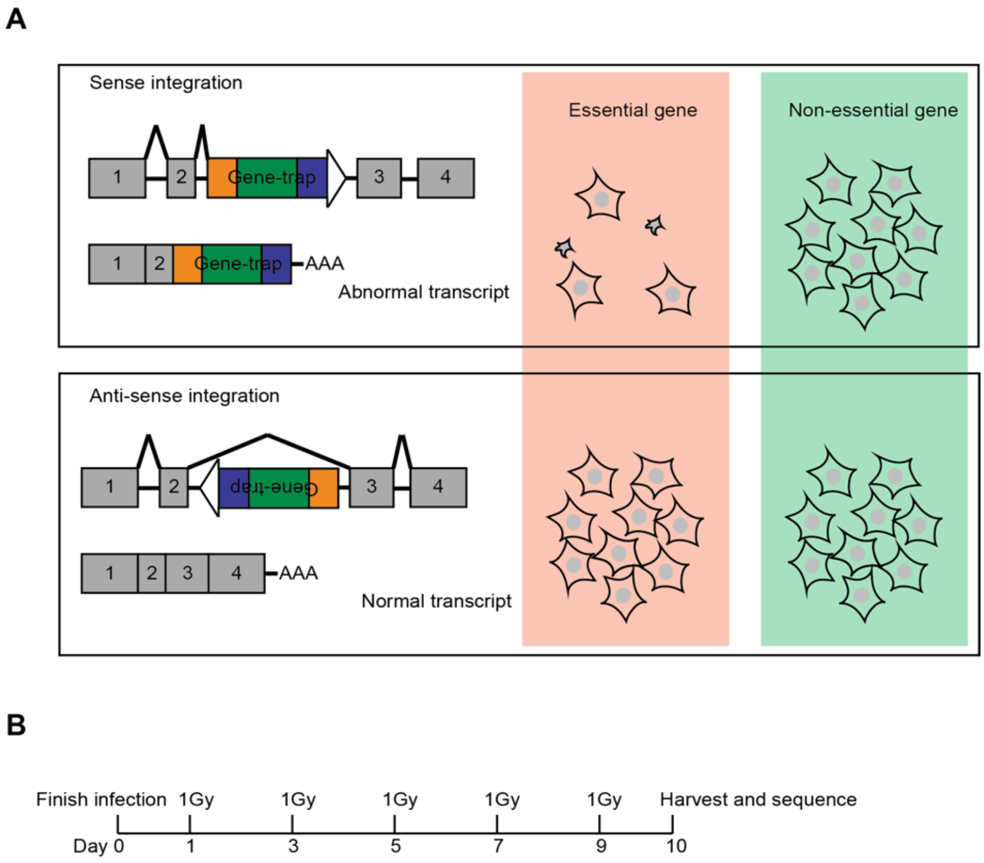


Figure S4

**Figure S4: A,** Schematic representation of the genetic screening approach in HAP1 cells. Random integration of the genetrap cassette can occur in sense or anti-sense orientation which generates an abnormal transcript (sense) or does not affect the transcript (anti-sense). Cells with sense integration of the cassette in a gene essential for survival under the culture conditions will be selectively depleted from the population. By calculating the number of sense vs total (sense + anti-sense) integrations per gene an ‘essentiality’ score for each gene is determined. **B,** Schematic representation of the experimental set-up. HAP1 cells infected with the genetrap virus were cultured for 10 days, while 1Gy irradiation was applied every other day. Genetrap insertions per gene were quantified and the percentage of sense-oriented integrations was compared with the percentage of sense integrations in untreated cells for each gene.

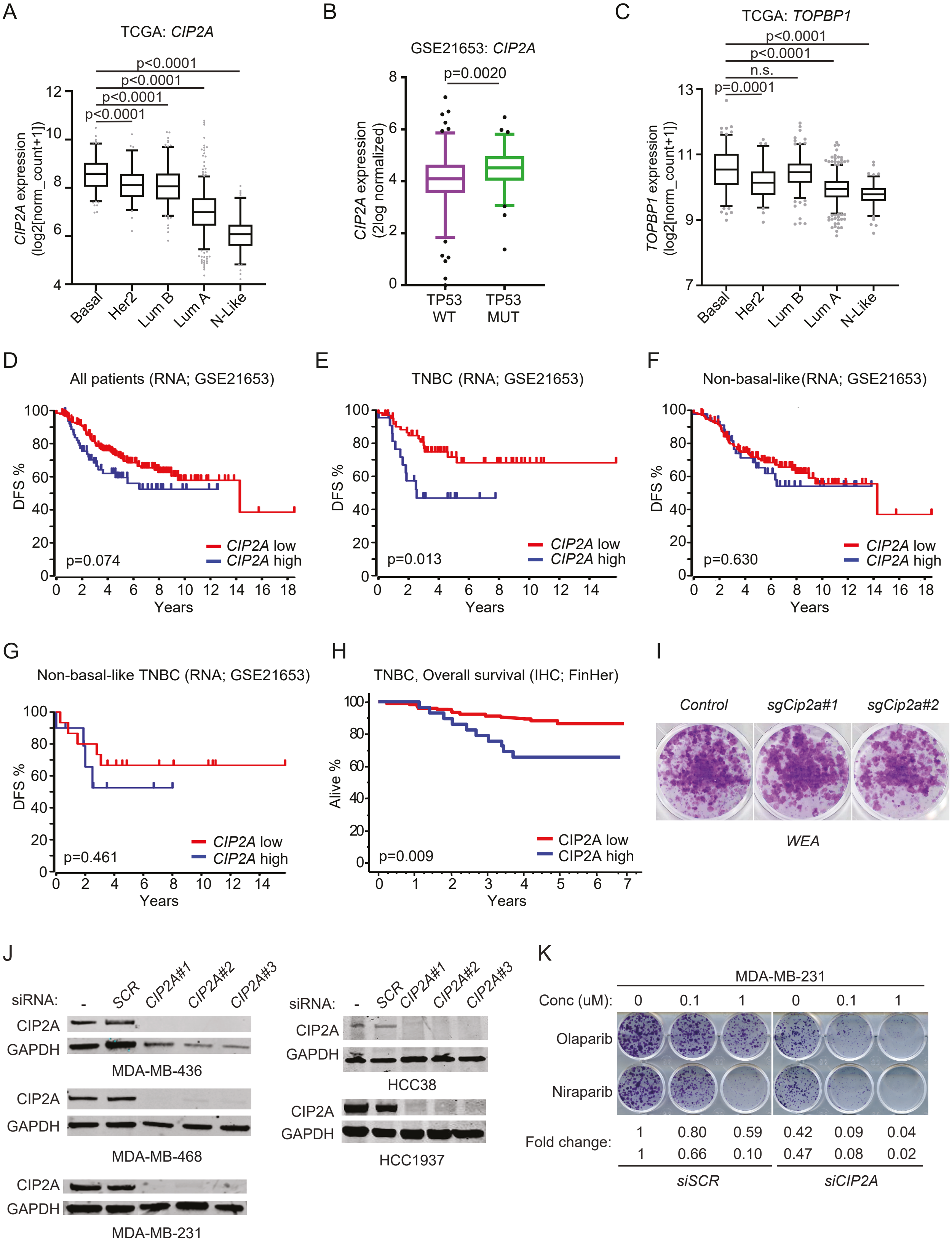
A
TCGA: CIP2A
p<0.0001
p<0.0001
p<0.0001
p<0.0001
CIP2A expression (log2[norm_count+1])
Basal
Her2
Lum B
Lum A
N-Like
B
GSE21653: CIP2A
p=0.0020
CIP2A expression (2log normalized)
TP53 WT
TP53 MUT
C
TCGA: TOPBP1
p<0.0001
p<0.0001
n.s.
p=0.0001
TOPBP1 expression (log2[norm_count+1])
D
All patients (RNA; GSE21653)
DFS %
p=0.074
CIP2A low
CIP2A high
Years
E
TNBC (RNA; GSE21653)
p=0.013
F
Non-basal-like (RNA; GSE21653)
p=0.630
G
Non-basal-like TNBC (RNA; GSE21653)
p=0.461
H
TNBC, Overall survival (IHC; FinHer)
Alive %
p=0.009
I
Control
sgCip2a#1
sgCip2a#2
WEA
J
siRNA:
-
SCR
CIP2A#1
CIP2A#2
CIP2A#3
CIP2A
GAPDH
MDA-MB-436
MDA-MB-468
MDA-MB-231
HCC38
HCC1937
K
MDA-MB-231
Conc (uM): 0 0.1 1 0 0.1 1
Olaparib
Niraparib
Fold change:
1 0.80 0.59 0.42 0.09 0.04
1 0.66 0.10 0.47 0.08 0.02
siSCR
siCIP2A

Figure S5

**Figure S5: A,** Expression of *CIP2A* in molecular breast cancer subtypes. **B,** Expression of *TOPBP1* in molecular breast cancer subtypes. **A, B,** Data derived from TCGA. Data is shown as 5-95 percentile boxplot with median and quartiles indicated. p-values by unpaired t-test. **C,** *CIP2A* expression in TP53 WT and TP53 mutated (MUT) breast cancers from GSE21653 cohort. Data is shown as 5-95 percentile boxplot with median and quartiles indicated. p-values by Mann-Whitney test. **D,** Disease-free survival (DFS) of *CIP2A* high (n=63) and *CIP2A* low (n=189) expressing from all breast cancer patients in GSE21653 cohort. **E,** Disease-free survival of *CIP2A* high (n=22) and *CIP2A* low (n=63) expressing TNBC breast cancer patients in GSE21653 cohort. **F,** Disease-free survival of *CIP2A* high (n=45) and *CIP2A* low (n=132) expressing non-basal-like (HER2+, luminal A, luminal B and normal-like) breast cancer patients in GSE21653 cohort. **G,** Disease-free survival of *CIP2A* high (n=10) and *CIP2A* low (n=15) expressing non-basal-like TNBC patients in GSE21653 cohort. **H,** Overall survival of CIP2A high (n=12) and CIP2A low (n=51) TNBC patients in FinHer cohort. **D to H**, p-values calculated by log rank test. **I,** Colony growth assays conducted on mammary tumor cell lines isolated from invasive lobular carcinoma-type (*WEA*: *E-Cadherin* mutant and oncogenic *Akt* expressing) mouse models; *Cip2a* was knocked out using CRISPR/Cas9 using 2 unique gRNAs. Shown are representative images of 2 biological replicates. **J,** Western blots confirming the downregulation of CIP2A using 3 unique *CIP2A* siRNAs used in Figure 5G. **K,** Colony growth assays in MDA-MB-231 cells treated with non-targeting (si*SCR*) and *CIP2A* targeting (siCIP2A) siRNA and treated with indicated concentrations of PARP inhibitors Olaparib and Niraparib. Relative colony areas are displayed under the representative images.

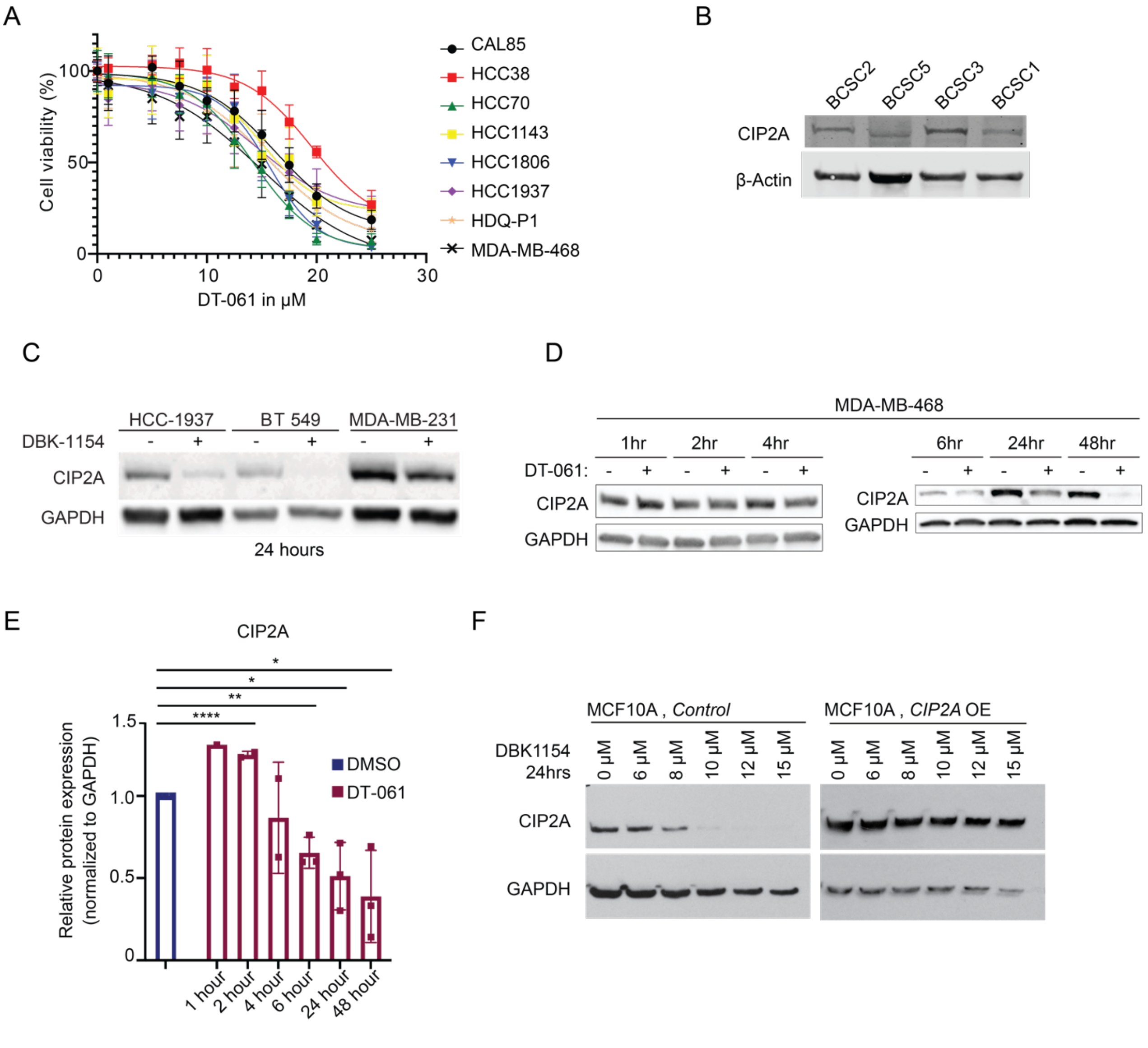


Figure S6

**Figure S6: A,** SMAP (DT-061) sensitivity profiles of eight BL-TNBC cell lines. Cell viabilities were measured using CellTiterGlo Luminescence Assay after 24 hrs of drug treatment. **B,** Western blot analysis of patient derived breast cancer stem-like cells (BCSCs) probed for CIP2A. β-Actin used as loading control. **C,** Western blots from HCC1937, BT549 and MDA-MB-231 cell lines treated with SMAP DBK-1154, 20μM for 24 hours indicating CIP2A protein downregulation after SMAP treatment. **D,** Time course of CIP2A protein expression in MDA-MB-468 cells treated with 20μM SMAP DT-061 for indicated time points **E,** Time course of CIP2A protein in MDA-MB-468 cells after treatment with 20μM DT-061 for the indicated time points. p-values calculated by unpaired t-test *p<0.05, **p<0.01, ****p<0.0001. **F,** Dose dependent effect of SMAP DBK-1154 on CIP2A protein expression in control vector overexpressing (MCF10A *Control*) and *CIP2A* overexpressing (*CIP2A* OE) MCF10A cells.

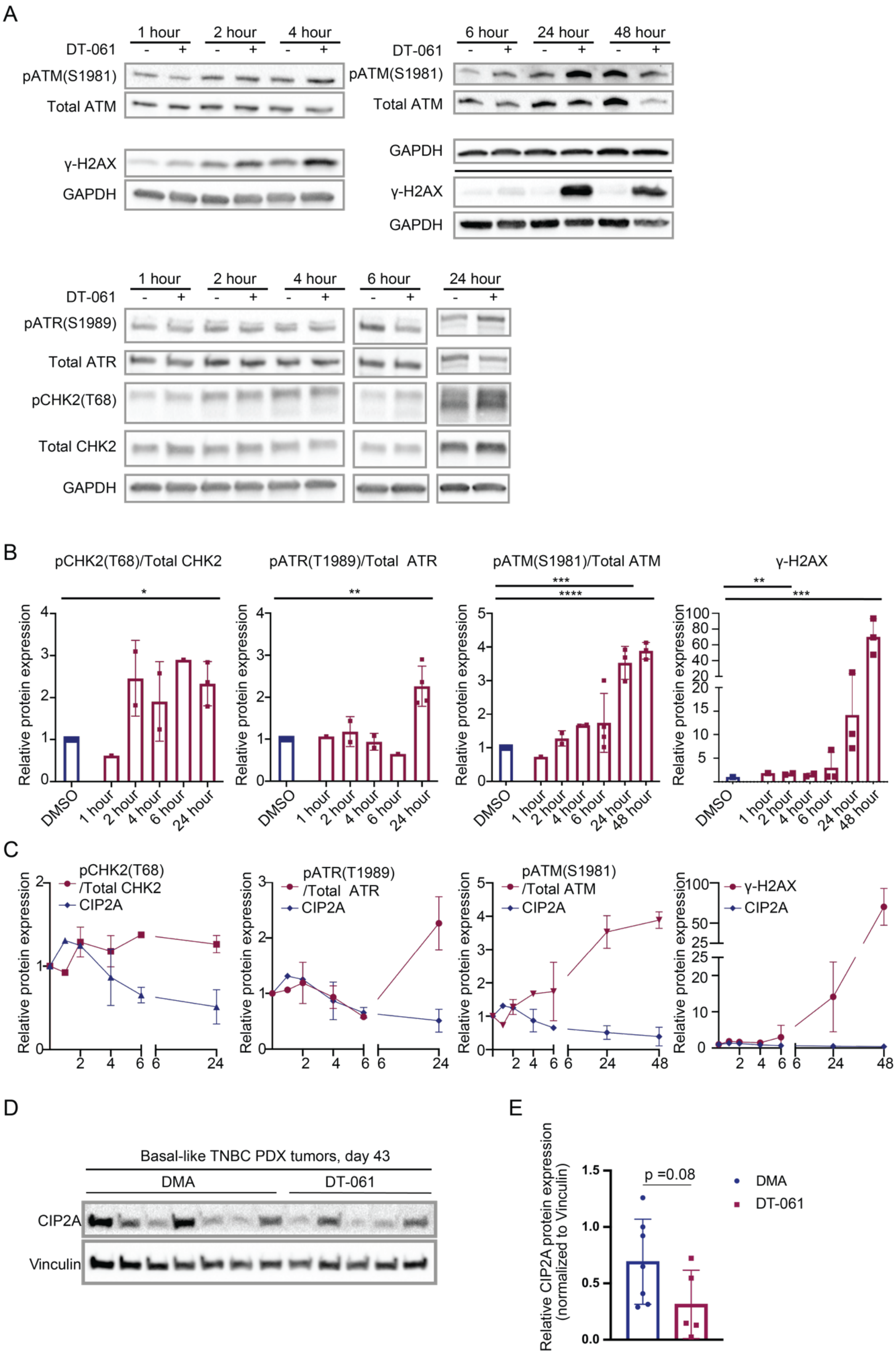
A
1 hour
2 hour
4 hour
6 hour
24 hour
48 hour
DT-061
pATM(S1981)
Total ATM
γ-H2AX
GAPDH
pATR(S1989)
Total ATR
pCHK2(T68)
Total CHK2
B
pCHK2(T68)/Total CHK2
pATR(T1989)/Total ATR
pATM(S1981)/Total ATM
γ-H2AX
Relative protein expression
DMSO
C
pCHK2(T68)
/Total CHK2
CIP2A
pATR(T1989)
/Total ATR
pATM(S1981)
/Total ATM
D
Basal-like TNBC PDX tumors, day 43
DMA
DT-061
CIP2A
Vinculin
E
Relative CIP2A protein expression
(normalized to Vinculin)
p =0.08
DMA
DT-061

Figure S7

**Figure S7: A,** MDA-MB-468 cells were treated with 20μM of SMAP (DT-061) and collected at 1, 2, 4, 6, 24 and 48 hours after treatment and assessed for markers of DNA Damage Response (DDR) by western blot **B,** Quantification of A representing n=2 at 1, 2 and 4 hours, and n=3 or 4 as indicated at 6, 24, 48 hours. **C,** Changes of DDR proteins relative to CIP2A expression over time. **D,** Western blots of representative tumor lysates from PDX model for the DMA treated mice (n=7) and SMAP DT-061 treated (n=5) mice probed for CIP2A. Vinculin is used as loading control. **E,** Quantifications of CIP2A levels from western blots in D. **B, C, and E,** Quantifications are represented as mean ± SD and p-values were calculated using unpaired t-test, *p<0.05, **p<0.01, *** p <0.001, ****p<0.0001.

#### Supplementary Materials and Methods:

<u>Animal experiments</u>

Development and genotyping of *Cip2a* genetrap hypomorphic mutant mouse has been previously described (1). Homozygous mice for genetrap cassette and wild type mice are referred here as *Cip2a-/-* and *WT* respectively. The *Cip2a-/-* mice were genotyped by PCR analysis of genomic DNA, which was further validated by assessing *Cip2a* mRNA by qRT-PCR from tissues collected upon autopsy.

To study role of CIP2A in skin carcinogenesis, DMBA/TPA protocol was used. The dorsal skin of mice was shaved a day before starting treatment. 50μg of DMBA dissolved in 200μl of acetone was administered once topically followed by 10μg of TPA dissolved in 150μl of acetone topically 10 days after the DMBA treatment for three times a week until the end of experiment (2). Mice were observed for formation of skin papillomas and the formed papillomas were counted weekly.

To study CIP2A in ovarian tumors *Cip2a+/-* mice were crossed with an ovarian cancer mouse model *TgMISIIR-Tag (3)*. In order to observe metabolic active tumor volume

(MATV) of ovarian tumors, female *WT;TgMISIIR-Tag* and *Cip2a-/-;TgMISIIR-Tag* mice were imaged with positron emission tomography (PET) (4). Mice anaesthetized with isoflurane were intravenously injected via the tail vein with 4.84 ± 0.43 MBq of 2-deoxy-2-[$^{18}$F]fluoro-D-glucose ([$^{18}$F] FDG) 120 min prior to a 20-min static scan (Inveon, Siemens). Volumes of interest were defined to obtain the MATV using a fixed (2.5% of injected dose per ml) value as a threshold value, which covered the entire tumor in all cases.

Mammary gland whole mount staining procedure was performed as described in (5). Briefly, mammary glands were isolated from *WT* and *Cip2a-/-* mice, spread onto super frost glass slides and left to air dry for 5 minutes. The tissue was then fixed with Carnoy's medium (60% of 95% Ethanol, 30% of Chloroform and 10% glacial acetic acid) for 4 hours at room temperature. The slides were rehydrated using decreasing ethanol series, stained for 2-3 days using Carmine alum staining solution (0.2% carmine and 5% Aluminum potassium sulphate dodecahydrate) and dehydrated using increasing ethanol series, followed by bleaching in xylene for 1-3 days at room temperature. The tissues were mounted with Permount and imaged using Zeiss SteREO Lumar.V12 stereomicroscope and 0.8X NEOLumar S objective. Multiple images per gland were merged into a mosaic picture using Photoshop.

The patient derived xenograft (PDX) model BCM 3204 (TG8-2296A) was a gift from Keri Lab at the Case Western Reserve University, Cleveland, Ohio, USA. The PDX model was generated and characterized as described in (6). Basal TNBC xenograft was derived from the primary breast tumor, pre-treatment. The tumor was positive for cytokeratin 19, cytokeratin 5/6, P53 and EGFR in both the patient biopsy and in consequent xenografts derived from the initial tumor sample. The patient was treated with a combination of doxorubicin (Adriamycin) and cyclophosphamide and was resistant to treatment.

PDX tumor fragments were implanted into the right inguinal mammary glands of adult (2 to 4 months old) female NOD/scid/γ mice. Tumors were measured every other day by caliper and body weight measured every four days. Mice were treated with SMAPs when average tumor volume reached about 100mm3 and were treated until mice had a body conditioning score of 1, when study had reached 43 days. At termination of study, mice received a final treatment 2 hours before sacrifice. Tumor tissue was collected and both formalin-fixed, for IHC, and snap frozen in liquid nitrogen for immunoblotting and mRNA analysis. SMAPs were delivered by oral gavage, twice a day (BID) at 5 or 15 mg/kg (mpk) in a solution of 10% N, N Dimethylacetamid (DMA) and 10% Solutol® HS15 (Kolliphor® HS 15) in sterile water. The *in vivo* PDX model was performed with approval from the Institutional Animal Care and Use Committee at Case Western Reserve University, which is certified by the American Association of Accreditation for Laboratory Animal Care under protocol # 2013-0132.

Analysis of human breast cancer patient sample cohorts

CIP2A was immunostained in FinHer breast cancer cohort as described previously (7). Tumors were classified as basal-like TNBC as described in (8). Shortly, tumors immunostained negative for ER, PR and HER2 and positive for EGFR and/or positive for the basal cytokeratin CK5 were classified as basal-like TNBC. Tumors immunostained negative for EGFR and CK5 and negative for ER, PR and HER2 were classified as non-basal-like TNBC. Tumors were immunostained for ER, PR, HER2, CK5 as previously described (8) and for EGFR by using EGFR pharmDx staining kit (Dako) according to manufacturer's instructions. FinHer study (HUCH 426/E6/00) was approved by an ethics committee of the Helsinki University Central Hospital (Helsinki, Finland).

The role of *CIP2A* in disease-free survival of breast cancer patients in the GSE21653 cohort was analyzed by using an online platform 'R2: Genomics Analysis and Visualization Platform' (https://hgserver1.amc.nl/cgi-bin/r2/main.cgi). Briefly, in previously published GSE21653 cohort (9) gene expression was analyzed using DNA microarrays from pre-treatment samples from 2145 invasive early breast adenocarcinomas in April 2019. Breast cancer subtype classification was performed by Single Sample Predictor classifier based on a list of 306 genes (10). Tumors immunostained negative for ER, PR and HER2 were classified as TNBC. In disease-free survival analysis *CIP2A* high group included cases with highest quartile of *CIP2A* expression. *CIP2A* low group included the rest of the cases in each analyzed breast cancer group. *CIP2A* expression was analyzed in TP53 mutant and WT cases from GSE21653 cohort. Tumors' TP53 mutation status was previously defined by TP53 immunohistochemistry in the cohort (9). Data was downloaded from 'R2: Genomics Analysis and Visualization Platform' in October 2019.

*CIP2A, TOPBP1* and *POLQ* gene expressions in different breast cancer subtypes in TCGA (The Cancer Genome Atlas Program) Breast Cancer data (Illumina HiSeq2000

RNA sequencing, n=1218) was obtained via UCSC Xena platform (https://xena.ucsc.edu/) (11) in June 2020. In order to study gene expressions in different molecular subtypes and in TNBC the following phenotypic identifiers were used: PAM50Call_RNAseq, ER_Status_nature2012, PR_Status_nature2012, and HER2_Final_Status_nature2012.

Haploid genetic screen

A haploid genetic screen to identify genes required for cell viability after treatment with IR was performed as described before (12). In short, genetrap retrovirus to mutagenize HAP1 cells was produced in HEK293T cells by transfection of the previously described genetrap plasmid (13) together with the packaging plasmids Gag-pol, VSVg and pAdv. At 48h after transfection, medium was harvested and concentrated by ultracentrifugation at 21.000 rpm for 2h at 4°C. Medium with retrovirus was collected and concentrated twice a day for three consecutive days. 40 million HAP1 cells were seeded per T175 flask and transduced with concentrated genetrap retrovirus in presence of 8ug/ml protamine sulfate (Sigma). The mutagenized HAP1 cells were expanded for 10 days while they were passaged or irradiated with 1Gy on alternating days. At day 11, cells were harvested, and pellets were fixed using BD fix buffer 1 (BD biosciences). To exclude diploid cells with potential heterozygous mutations, cells were stained using DAPI in PBS + 10% FCS and sorted on a MoFlo Astrios Cell Sorter (Beckman Coulter). Approximately 30 million G1 (1n) cells were sorted of which DNA was collected using a Qiagen DNA mini kit. De-crosslinking was achieved by overnight incubation in lysis buffer (buffer AL, Qiagen) and Proteinase K (Qiagen) at 56°C.

A linear amplification polymerase chain reaction (LAM-PCR) was performed on the total genomic DNA isolated from 30 million cells to amplify insertion sites. The reaction was performed in 120 cycles with alternating annealing temperature of 58°C and extension temperature of 68°C using a double-biotinylated primer. Biotinylated single-stranded DNA (ssDNA) products were obtained by magnet after 2h incubation with M270 streptavidin-coated magnetic beads (Life Technologies). A ssDNA linker was ligated to the biotinylated products followed by a PCR reaction to introduce adaptor sequences needed for Illumina HiSeq2000 or HiSeq2500 (Illumina). The obtained reads were aligned to the human genome (GRCh38) using Bowtie-align-s version 1.2.2 to define insertion sites. The fraction of sense integrations was determined after mapping of sense and antisense integrations for each gene. The fraction of sense integrations after IR treatment was compared with 4 control screens reporting gene-essentiality for survival in untreated conditions (12). Binomial testing with FDR correction (Benjamini Hochberg) was used to identify genes with a significant drop in sense-insertions after the IR treatment applied here.

Exome-sequencing and mutation load analysis

For exome-sequencing genomic DNA was isolated from control and DMBA-treated *WT* and *Cip2a-/-* mouse mammary glands by using DNeasy Blood & Tissue Kit (Qiagen). Samples were sequenced in a lane of PE 100bp HPM on HiSeq2500 (Illumina) to reach 50x coverage. Exome-sequencing was performed at Netherlands Cancer Institute (Amsterdam, Netherlands). The quality control of the sequenced reads was performed with FastQC (14), with all the samples passing the quality control. The reads were aligned to GRCm38 reference genome using Burrows-Wheeler Aligner (BWA)-MEM v0.7.17 software (15). The subsequent steps, including the pre-processing of the aligned files and the variant calling, were performed with Genome Analysis Toolkit GATK v3.7.0 following the Best Practices guidelines (16). For variant calling GATK's HaplotypeCaller was used. All the single nucleotide variants as well as short indels within the expressed genes were included in the inspection of mutation load.

RNA isolation and qRT-PCR

RNA was isolated using NucleoSpin RNA kit (Macherey-Nagel) with DNase I treatment. RNA was converted to cDNA using Random Primers, Recombinant RNasin ribonuclease inhibitor, M-MLV RT RNase (H-) point mutant (Promega) and dNTP mix (Thermo Fisher). cDNA was analysed by using primers listed in Table S4 using QuantStudio™ 12K Flex Real-Time PCR System (Thermo Fisher Scientific). *Actb* and/or *Gapdh/GAPDH* were used as reference genes.

RNA-sequencing and Gene Set Enrichment Analysis

RNA was isolated from HCC38 cells transfected with indicated siRNAs after 72 hours as mentioned in previous sections. *CIP2A* knockdown was validated by using qRT-PCR using Primers listed in Table S4. Human *GAPDH* was used as a reference gene. The samples were submitted for sequencing after the *CIP2A* downregulation was confirmed. The sequencing was performed at Finnish Functional Genomics Centre (Turku Bioscience Centre, Turku, Finland). The samples were sequenced with the Illumina HiSeq2500 instrument using single-end sequencing chemistry with 50 bp read length. A ranked list was created from the differentially regulated genes, and GSEAPreranked analysis was run with default settings. Hallmarks dataset was used to generate the enriched gene sets.

Drug screening in breast cancer stem-like cells

The drug sensitivity and resistance testing experimental design was adopted from (17). Briefly, thirteen compounds (3 SMAPs & 10 chemotherapies, mentioned in Table S5) were plated to white clear bottom 384-well plates (Corning #3712) in 5 increasing concentrations in 10-fold dilution steps covering a 10,000-fold concentration range using an Echo 550 Liquid Handler (Labcyte). 100μM benzethonium chloride (BzCl2) was used as positive control whereas 0.1% dimethyl sulfoxide (DMSO) was used as negative control. All subsequent liquid handling was performed using MultiFlo FX multi-mode dispenser (BioTek). The pre-dispensed compounds were dissolved in 5 μl of culture media with CellTox Green (1:2000 final volume, Promega) and left in a plate shaker at room temperature for 30 min. 20μl cell suspension was dispensed into the drug plates. After 72 hours incubation, fluorescence (cytotoxicity) was measured using PheraStar plate reader (BMG Labtech). The raw fluorescence data were analyzed in Breeze software (18) using an inhouse developed data analysis pipeline at Institute for molecular Medicine Finland (FIMM), to calculate the drug sensitivity scores (DSS)(19).

Cloning

Lentiviral plasmid pWPI was purchased from Addgene. pWPI-CIP2A-V5: Human CIP2A was cloned into pWPI vector using Phusion Green Hot Start II High-Fidelity PCR Master

Mix (Thermo Fisher Scientific, F566S). The forward primer containing restriction sites for SwaI and PacI respectively were used and reverse primer contained V5 tag. As a DNA template, pcDNA3.1_kozak_CIP2A_1-905_V5His was used. PCR product and pWPI were digested using Fast Digest SmiI (FD1244, SwaI isoschizomer) and PacI (FD2204, both from Thermo Fisher Scientific), with slightly modified manufacturer´s protocol to allow maximum digestion efficiency. The products were ligated using 2.5 units of T4 DNA Ligase (EL0011 5U/ µL, from Thermo Fisher Scientific) and approximately 3-fold mass excess of insert over vector. For transformations and plasmid propagations, DH5α cells were used throughout. The integrity of the ligation was verified by PCR and by sequencing conducted at FIMM Helsinki.

Flow cytometry

*Characterization of MMECs:* For characterizing and quantifying the proportion of basal and luminal cells from the *WT* and *Cip2a-/-* MMECs, flow cytometry protocol described previously was used (5). Briefly, cells were suspended in Tyrodes buffer and approximately 0.5 million cells were used per labeling. Fluorophore-conjugated antibodies (dilutions in Tyrodes buffer described in Table S4) were incubated for 30 minutes at 4°C, washed twice in Tyrodes buffer, and fixed with 4% PFA for 10 min at room temperature. Samples were analyzed using BD LSR Fortessa flow cytometer. Labeling was done with two antibody pairs (CD24/CD29 or CD24/CD49f).

*Gating strategy:* First, the single and live cells were gated using FSC/SSC. From live cells, the lineage negative ($CD31^{neg}$ and $CD45^{neg}$) and the $CD24^{pos}$ cells (epithelial) were gated for further analysis. Within the epithelial cell population, the basal cells ($CD24^{low}$ and $CD29^{high}$ or $CD49f^{high}$) MMECs and luminal ($CD24^{high}$ and $CD29^{low\text{-}neg}$ or $CD49f^{low\text{-}neg}$) MMECs were gated. The proportions of luminal and basal cells were quantified using FlowJo software.

*Sorting luminal and basal mouse mammary epithelial cells (MMECs):* Luminal and basal mouse mammary epithelial cells were sorted by flow cytometry from adult *WT* female mice according to previously published protocol (20). Shortly, mammary gland tissue was mechanically dissociated and treated for two hours with collagenase and hyaluronidase (StemCell Technologies). Red blood cells were lysed with ACK buffer followed by 5 min treatment with prewarmed Trypsin-EDTA, 2 min treatment with prewarmed dispase (StemCell Technologies) with DNase I (Sigma) and filtration through 70µm mesh and washing with 2% FCS in HBSS. Single cells were blocked with rat IgG (Sigma) 10mg/ml 1:1000 in 2% FCS in HBSS. Cells were stained with antibody mix on ice for 45 min. The used antibodies are included in the Table S4. Flow cytometry was performed with BD FACSAria IIIu (Becton Dickinson Biosciences) by using Diva software. First, single cells were gated using FSC/SSC followed by gating live cells by using 7-AAD staining. Next, lineage negative (CD45, TER119 and CD31 negative) cells were gated out to analyse basal and luminal epithelial cells. Luminal cell population was $CD24^{high}$ and $CD49f^{low\text{-}neg}$ and basal cell population was $CD24^{low}$ and $CD49f^{high}$.

Proximity ligation assay (PLA)

The PLA assay was performed according to the manufacturer's protocol (Duolink® PLA, Sigma). Briefly, HeLa cells were plated on coverslips and 24 hours later, the cells were fixed with 4% paraformaldehyde 15 minutes at room temperature, and then cells were permeabilized with 0.5% Triton X-100 in PBS on ice for 5 minutes. Next, cells were blocked with blocking solution, and incubated in a pre-heated humidity chamber for 30 min at 37°C, followed by incubating the primary antibodies in blocking buffer (details of PLA antibodies used mentioned in Table S4) overnight at 4°C. The cells were washed with buffer A, and the PLA probe was incubated in a pre-heated humidity chamber for 1 hour at 37°C, followed by ligase reaction in a pre-heated humidity chamber for 30 minutes at 37°C. Next, amplification polymerase solution for PLA was added, followed by incubating the cells in a pre-heated humidity chamber for 100 min at 37°C. After amplification, the coverslips were washed with buffer B, and mounted with DAPI. PLA signal was detected by using a confocal microscope (LSM780, Zeiss). The images were quantified using Cell Profiler.

Data mining from public datasets

For co-expression analyses, Broad Institute's Cancer Cell Line Encyclopedia (CCLE) dataset containing 1739 samples (out of which 1156 had mRNA expression information) was used and analyzed using cBioportal website (http://www.cbioportal.org/). For analysis of genome wide gene dependencies, Broad Institute's DepMap portal (https://depmap.org/portal/) was used. Data was extracted from Achilles project CRISPR screens comprising of 739 cell lines (Avana 2020Q1 release). *CIP2A* gene dependence across all 33 breast cancer cell lines was sorted using gene dependency CERES score provided by the screen. CERES score is a gene dependency score generated taking into account copy number of the gene and also the guide RNA activity score (21). The score of -1 is the median of all essential genes and 0 is the score of a non-essential gene. Lower CERES score indicates that the cell line is more likely to be dependent on that particular gene.

Yeast two-hybrid screen

The yeast two-hybrid screen was performed by Hybrigenics. The full-length *CIP2A* was used as a bait, and the library in the screen was breast tumor epithelial cells (T47D, MDA-MB-468, MCF7, BT20).
